# Neurophysiological measures of covert semantic processing in neurotypical adolescents actively ignoring spoken sentence inputs: A high-density event-related potential (ERP) study

**DOI:** 10.1101/2024.02.29.582736

**Authors:** Kathryn K. Toffolo, Edward G. Freedman, John J. Foxe

**Affiliations:** The Frederick J. and Marion A. Schindler Cognitive Neurophysiology Laboratory, The Del Monte Institute for Neuroscience, Department of Neuroscience, University of Rochester School of Medicine and Dentistry, Rochester, New York, USA 14620

**Keywords:** EEG, N400, P600, Late Positive Component, Passive, Recognition Potential

## Abstract

Language comprehension requires semantic processing of individual words and their context within a sentence. Well-characterized event-related potential (ERP) components (the N400 and late positivity component (LPC/P600)) provide neuromarkers of semantic processing, and are robustly evoked when semantic errors are introduced into sentences. These measures are useful for evaluating semantic processing in clinical populations, but it is not known whether they can be evoked in more severe neurodevelopmental disorders where explicit attention to the sentence inputs cannot be objectively assessed (i.e., when sentences are passively listened to). We evaluated whether N400 and LPC/P600 could be detected in adolescents who were explicitly ignoring sentence inputs. Specifically, it was asked whether explicit attention to spoken inputs was required for semantic processing, or if a degree of automatic processing occurs when the focus of attention is directed elsewhere? High-density ERPs were acquired from twenty-two adolescents (7-12 years), under two experimental conditions: 1) individuals actively determined whether the final word in a sentence was congruent or incongruent with sentence context, or 2) passively listened to background sentences while watching a video. When sentences were ignored, N400 and LPC/P600 were robustly evoked to semantic errors, albeit with reduced amplitudes and protracted/delayed latencies. Statistically distinct topographic distributions during passive versus active paradigms pointed to distinct generator configurations for semantic processing as a function of attention. Covert semantic processing continues in neurotypical adolescents when explicit attention is withdrawn from sentence inputs. As such, this approach could be used to objectively investigate semantic processing in populations with communication deficits.

## INTRODUCTION

Individual words, prosodic cues, and the ongoing context of a sentence are used to comprehend meaning within spoken language. For example, if one were to hear the sentence *“I baked a birthday clue”*, most native English speakers would reasonably expect the ending to be *‘cake’*, which is semantically congruous with the prior context. However, in this instance, the highly improbable word *‘clue’* is semantically incongruous, and clearly violates the sentence context. Differences in the neurophysiological responses to manipulations of semantic congruency such as this can be readily measured using scalp-recorded event related potentials (ERPs). A series of three well-characterized ERP response modulations have been described in the literature that have differing amplitudes depending on these types of manipulations of semantic congruence: 1) the recognition potential (RP), recorded over left frontal scalp (∼250 ms); 2) the N400 component, recorded over midline central and parietal scalp (∼400 ms); and 3) the late positive component (LPC/P600), recorded over left parietal scalp (∼600 ms) (Toffolo et al., 2022).

Literature about the RP is predominated by visual word/picture paradigms for which the RP is generally found over the visual word form area (Dien et al., 2003; Martín-Loeches, 2007; Martín-Loeches et al., 1999). However, in a recent study by our group using spoken sentence stimuli, an RP-like component was recorded over left frontal scalp (∼250 ms), which suggests that there are scalp distribution differences based on the sensory modality of the stimulation (Toffolo et al., 2022). The RP is typically larger in amplitude in response to congruent stimuli and as such, is thought to represent the point at which an individual recognizes the target word following congruent context (Fernandez and Smith Cairns, 2011; Martín-Loeches et al., 2004; Proverbio and Riva, 2009). The N400 component, which has been much more extensively studied, is a broad negative deflection that peaks approximately 400 ms after the presentation of a stimulus that is related (congruent) or unrelated (incongruent) to the prior context (Block and Baldwin, 2010; Kutas and Hillyard, 1984, 1980; Lau et al., 2008; Luck, 2005; Osterhout and Holcomb, 1992; Toffolo et al., 2022). Because stimuli incongruent to the context result in larger N400 amplitudes (e.g., *‘clue’* in the example given above), the N400 is considered an index of semantic integration (Brouwer et al., 2017; Colin Brown and Hagoort, 1993; Connolly and Phillips, 1994; Hagoort, 2008; Kaan, 2007; Kutas and Hillyard, 1980; Lau et al., 2008; Osterhout and Holcomb, 1992; Toffolo et al., 2022). Lastly, the LPC/P600 is a component that has more positive amplitudes (∼600 ms) in response to stimuli that are incongruent to the previous context (Brouwer et al., 2017; Gouvea et al., 2010; Kaan et al., 2000; Van Herten et al., 2005; Wang et al., 2009). Given its later temporal course, following the N400, it is thought to reflect the reanalysis of semantic information and/or the integration of semantic and syntactic information (Ainsworth-Darnell et al., 1998; Braeutigam et al., 2008; Brouwer et al., 2017; Brouwer and Hoeks, 2013; Fitz and Chang, 2019; Friederici, 2011, 2002; Gouvea et al., 2010; Kim and Osterhout, 2005; Osterhout and Holcomb, 1992; Schacht et al., 2014; Toffolo et al., 2022; Van Herten et al., 2005; Wang et al., 2009).

Research using these dependent measures often focuses on semantic comprehension in individuals actively paying attention to, and acting upon, the stimuli. However, many populations of key interest to clinical researchers are limited to passive observation or listening paradigms due to impairments in communication, such as those with disorders of consciousness (DOC) and intellectual or developmental disabilities (IDD: e.g. Rett Syndrome, minimally-verbal autism spectrum disorder (mvASD)). Because instruction often cannot reasonably be given to these individuals and attentional engagement cannot be objectively monitored, clinical assessment and behavioral observation are primarily used for diagnosis and treatment. Although behavioral measures are important, neurophysiological methods provide a potentially objective insight on covert sensory-perceptual and cognitive abilities in these clinically severe populations (Brima et al., 2024b, 2024a, 2019; Foxe et al., 2016; Giacino et al., 2014; Knight et al., 2020; Rohaut et al., 2015; Rohaut and Naccache, 2018; Seel et al., 2010). Simple error prediction paradigms have previously been used to characterize lower level auditory sensory-perceptual processing in individuals with mvASD and to predict better outcomes in patients with DOC (Giacino et al., 2014; Knight et al., 2020; Risetti et al., 2013; Rohaut et al., 2015). Alternatively, assessment of higher-order semantic processing abilities via the N400 component has led to mixed results in the same populations, especially at the single participant level, generating a need for development of better tools for measuring semantic comprehension levels in populations with these sorts of communication impairments (Cantiani et al., 2016; Daltrozzo, 2009; Kotchoubey and Daltrozzo, 2005; Pajankar, 2022; Rohaut et al., 2015).

A considerable body of work has addressed the role of active attentional engagement with semantic inputs on the evocation of the N400, with generally consistent findings showing N400 amplitude attenuations when the input stimuli are not actively attended. For example, when participants were asked to listen passively (i.e. no active response was required) to congruent/incongruent stimuli, the N400 and a related priming effect were elicited, but with attenuated amplitudes relative to an active engagement paradigm (Cantiani et al., 2016; Erlbeck et al., 2014). Moreover, when participants were asked to completely ignore the stimuli and watch a film, N400 amplitudes were not only reduced, but the semantic priming effect was strongly attenuated or eliminated (Erlbeck et al., 2014; Relander et al., 2009). Similarly, only stimuli in the attended stream during presentation of two concurrent streams elicited an N400 and semantic priming effect, and when attention was divided between two streams, the N400 effect was reduced (Bentin et al., 1995; Hohlfeld et al., 2015; Hohlfeld and Sommer, 2005; Hubbard and Federmeier, 2021; McCarthy and Nobre, 1993; Okita and Jibu, 1998). Interestingly, when a participant’s attention was focused on unrelated or non-semantic elements, N400 amplitudes were reduced and the priming effect was attenuated or eliminated, but when attention was still focused on word meaning (i.e. distinguishing words from non-words), an N400 was still evoked in the ostensibly unattended stream (Chwilla, 2022; Chwilla et al., 1995; Erlbeck et al., 2014; Hohlfeld and Sommer, 2005; Maxfield, 1997; Tripier, 2018). The N400 in response to congruency differences has even been reported in individuals who are sleeping (Bastuji et al., 2002; Brualla et al., 1998; Ibáñez et al., 2006, 2009; Perrin et al., 2002). There has been minimal research of the effects of attention on the RP or the LPC/P600, but some studies have suggested that the amplitudes of these components are also substantially attenuated when attention is withdrawn (Chwilla, 2022; Contier et al., 2022; Rudell and Hua, 1996). In summary, although the degree of attenuation differs across studies and across the considerable variations in paradigm used, reductions in explicit attention to semantic inputs have relatively consistently led to amplitude reductions in the N400, as well as reductions in RP and LPC/P600 amplitudes.

The current work was motivated by a number of factors. First, there has been little consistency in the nature of the paradigms used to evoke these dependent measures and there has been very little standardization within the field. Here, the auditory stimuli were drawn from a publicly-available stimulus set of full sentences (with and without semantic ending errors) which was verified to evoke robust RP, N400, and LPC/P600 modulations in neurotypical adults (Toffolo et al., 2022). Second, our use of full sentence stimuli is distinct from much of the prior semantic attention literature, which has typically investigated paired-word comparisons (e.g. Sofa - Chair vs. Sofa - Airplane). It is not an unreasonable proposition that these non-ecological paired-word presentations evoke a distinct form of semantic comprehension that may not recapitulate the typical way in which humans process semantic inputs. Alternatively, full sentences provide more context, require that the listener maintains/utilizes semantic knowledge of multiple words, and as such, may target higher-level semantic comprehension abilities. Because a major purpose of this study was to develop a tool whereby higher-level semantic processing can be assessed without explicit attention to the inputs, we reasoned that providing the participant with more naturalistic sentential context might contribute more information about the extent to which semantic information can be processed covertly. Lastly, the great majority of studies investigating attentional influences on semantic processing have been conducted in adult observers. The current work aimed to develop a tool that could be deployed in pediatric clinical populations with communication difficulties, where explicit attention cannot be controlled or explicitly measured. To this end, 22 adolescents, between the ages of 12-17 years, were recruited to assay whether attention is needed to evoke these components for accurate measurement of semantic comprehension. As with our prior work, all stimuli, code and the 22 high-density EEG datasets are made publicly available to the community (Toffolo et al., 2022).

This study presents auditory semantic sentences to adolescent participants in both an active context with a direct behavioral component (i.e. responding to each stimulus) and a passive context (i.e. watching an unrelated silent television show). Overall, we hypothesized that, relative to the adults who experienced the same stimuli in our prior work (Toffolo et al., 2022), similar RP, N400, and LPC/P600 modulations would be evoked in adolescents during an active engagement paradigm where they were required to determine whether the last word of a sentence was congruent or incongruent to the prior context. Alternatively, because the passive paradigm of this study asks the participant to ignore the stimuli and only focus on a silent show of their choice (a feasible task design when working with populations with communication differences such as mvASD and Rett Syndrome), we hypothesized that the RP, N400, and LPC/P600 would continue to be detected, but that their amplitudes would be significantly reduced relative to the active engagement paradigm. We reasoned that if these components remained robustly detectable when the inputs were explicitly ignored, that the current paradigm could be assumed to have important utility in the objective measurement of semantic processing in pediatric populations with severe communication difficulties.

## EXPERIMENTAL PROCEDURES

### Participants

Twenty-two NT adolescents between the ages of 12-17 provided informed assent with parental permission to enroll in the study, and participated in both active and passive paradigms. One active paradigm dataset (sub-07) was removed from analysis due to excessive noise in the recordings. Final analyses were conducted on twenty-one adolescents in the active paradigm and twenty-two in the passive paradigm. Of these twenty-two participants, twelve were female and four were left-handed. Participants identified themselves as 54.5% (12/22) White, 18.2% (4/22) White and Hispanic or Latino, 13.6% (3/22) more-than-one race and Hispanic or Latino, 9.0% (2/22) more-than-one race, and 4.5% (1/22) Black or African American. No participant identified that their gender was different to their biological sex, so only sex was reported in participant information. All participants were required to have English as their first language, and eight reported being bi- or multi-lingual. Two participants were bilingual from birth. Demographic information for all participants is reported in **Table 1**.

**Table 1.**
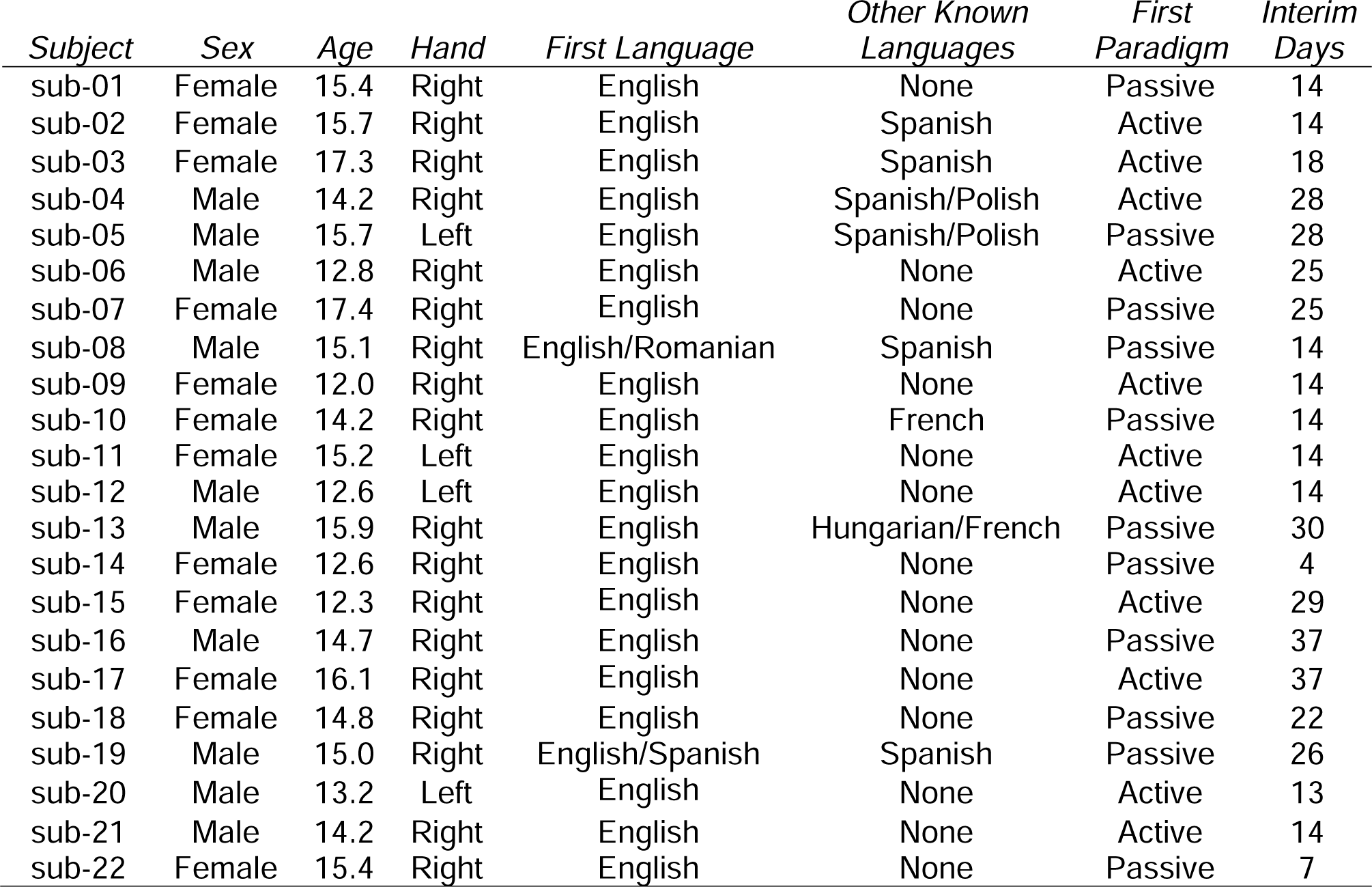
Demographic Data of the Participants. Included for each participant is their biological sex, age in years, handedness, first language, other fluent known languages, which paradigm was conducted first, and the number of intervening days between the first and second visit. Data from sub-07 were only included in the “passive” analyses.

| *Subject* | *Sex* | *Age* | *Hand* | *First Language* | *Other Known Languages* | *First Paradigm* | *Interim Days* |
| --- | --- | --- | --- | --- | --- | --- | --- |
| sub-01 | Female | 15.4 | Right | English | None | Passive | 14 |
| sub-02 | Female | 15.7 | Right | English | Spanish | Active | 14 |
| sub-03 | Female | 17.3 | Right | English | Spanish | Active | 18 |
| sub-04 | Male | 14.2 | Right | English | Spanish/Polish | Active | 28 |
| sub-05 | Male | 15.7 | Left | English | Spanish/Polish | Passive | 28 |
| sub-06 | Male | 12.8 | Right | English | None | Active | 25 |
| sub-07 | Female | 17.4 | Right | English | None | Passive | 25 |
| sub-08 | Male | 15.1 | Right | English/Romanian | Spanish | Passive | 14 |
| sub-09 | Female | 12.0 | Right | English | None | Active | 14 |
| sub-10 | Female | 14.2 | Right | English | French | Passive | 14 |
| sub-11 | Female | 15.2 | Left | English | None | Active | 14 |
| sub-12 | Male | 12.6 | Left | English | None | Active | 14 |
| sub-13 | Male | 15.9 | Right | English | Hungarian/French | Passive | 30 |
| sub-14 | Female | 12.6 | Right | English | None | Passive | 4 |
| sub-15 | Female | 12.3 | Right | English | None | Active | 29 |
| sub-16 | Male | 14.7 | Right | English | None | Passive | 37 |
| sub-17 | Female | 16.1 | Right | English | None | Active | 37 |
| sub-18 | Female | 14.8 | Right | English | None | Passive | 22 |
| sub-19 | Male | 15.0 | Right | English/Spanish | Spanish | Passive | 26 |
| sub-20 | Male | 13.2 | Left | English | None | Active | 13 |
| sub-21 | Male | 14.2 | Right | English | None | Active | 14 |
| sub-22 | Female | 15.4 | Right | English | None | Passive | 7 |

### Stimuli

Auditory sentences with and without semantic ending errors were acquired from a publicly available semantic stimulus set (https://doi.org/10.5061/dryad.9ghx3ffkg), and were described in detail previously (Toffolo et al., 2022). Of the 442 stimuli provided in this set, the current study used 404 exemplars (202 congruent and 202 incongruent sentence pairs). These non-prosodic sentences ranged from four to eight words in length, and have associated cloze probability scores (i.e. the likelihood that a given sentence-ending word would be provided by typical observers (Kutas and Hillyard, 1984)), and were designed for use with children 5 years and older.

### Experimental design

Experiments were conducted in a sound attenuating and electrically shielded booth (Industrial Acoustics Company, The Bronx, NY), fit with a computer monitor (Acer Predator Z35 Curved HD, Acer Inc.) and a standard keyboard (Dell Inc.). Participants wore BioSemi 128-electrode caps (BioSemi B.V. Amsterdam, the Netherlands) for EEG data collection. Both passive and active task paradigms were created with Presentation® Software (Version 18.0, Neurobehavioral Systems, Inc. Berkeley, CA). Half of the participants underwent the active task on their first visit and the passive task on their second visit. The remaining participants underwent the reverse. There was a predetermined minimum of four days between visits for any given participant (Range = 4-37 days; Average = 20 days). During the consenting process, the purpose of the experiment was explained to all participants. All participants were made aware that sentence stimuli, some of which would be nonsensical, would be presented to them on both days of the experiment, regardless of whether they began with the active or the passive task on Day 1. Individuals were asked to refrain from excessive movement during experimental sessions, until break periods. A total of 402 stimuli were presented to participants in a randomized order on both experimental visits. The active paradigm had an additional two sentences used for practice that were not included in analyses. These practice trials were the same for every participant and presented after verbal instruction.

For the active paradigm, individuals were asked to focus on a fixation cross throughout the task. Before the two practice trials, all instructions were explained both on the screen and through headphones (Sennheiser HD 600 electronic GmbH & Co. KG, USA). Corrective feedback was only given during practice trials and not experimental trials. During experimental trials, an auditory sentence was played through headphones while a fixation cross was on the screen. This was followed by a two second pause, which was in turn followed by a prompt asking if the sentence ended as expected (the prompt was presented both visually and auditorily). Participants would then respond with a left arrow key press if a sentence ended unexpectedly (incongruent) or a right arrow key press if it ended as expected (congruent). Between a response and the start of the next sentence stimulus was a two second pause. Participants were given optional breaks every 20 or 40 stimuli and could continue with the experiment by pressing the spacebar. For the passive paradigm, individuals were instructed to simply ignore the auditory sentence stimuli and watch a show of their choice without sound or subtitles. No response was required for this paradigm and between each sentence stimulus was a four second gap (i.e. from the end of the last word in the preceding sentence to the beginning of the first word in the next). No breaks were given since participants were watching a continuous television show. Stimulation in the passive paradigm lasted approximately 46 minutes.

### Behavioral analysis

Accuracy was calculated by dividing the number of correct responses by the total number of answered trials and multiplying by 100. The responses to all trials were included regardless of whether they were eliminated in EEG preprocessing. The d-prime and c-bias values were calculated by separating correct and incorrect responses by condition. Specifically, if a left arrow was pressed in responses to an incongruent sentence, it was considered a “hit”, while if a right arrow key was pressed, it was considered a “miss”. If a left arrow was pressed in response to a congruent sentence, it was considered a false alarm or “fa”, while if a right arrow key was pressed, it was considered a correct inclusion or “ci”. After these values were added, the following equations were applied: d-prime = z(hits/(hits+miss)) – z(fa/(fa+ci)); c-bias= z(hits/(hits+miss)) + z(fa/(fa+ci)) (Green and Swets, 1966).

### Data preprocessing

Data were digitized online at a rate of 512Hz, DC to 150 Hz pass-band, and referenced to the common mode sense (CMS) active electrode. EEG data were then preprocessed offline over three preprocessing stages via custom-written in-house scripts utilizing EEGLAB functions (Delorme and Makeig, 2004). The purpose of the first preprocessing stage was to identify what channels and sections of the continuous data contained significant artifact/movement. The second preprocessing stage applied Independent Component Analysis (ICA) to the data. When used for EEG analysis, ICA is a technique that separates out artifacts embedded in the EEG recording from brain data (Delorme and Makeig, 2004). The sole purpose of this second preprocessing stage was to acquire component weights (i.e. likelihoods) of the data for brain, muscle, eye, heart rate, line noise, channel noise, or other artifacts. The final preprocessing stage rejected the bad channels/data identified in the first stage, then applied the component weights from the second stage to the filtered data. After subsequently marking components that contained artifacts for rejection, this final preprocessing stage resulted in the filtered data used for analysis.

During the first preprocessing stage, channels with flat-line data longer than 5 sec were removed prior to filtering. A Chebyshev II spectral filter with a pass band of 0.1 - 40 Hz was used to filter the data before they were inspected for channel rejection. There were two steps to the channel rejection process. During the first, channels were marked for rejection by two automatic methods: 1. Through “findbadchans”, which marked channels that exceeded more than three standard deviations beyond the mean variance and amplitude from all neighboring electrodes, and 2. Through functions within EEGLAB’s “clean_channels”, which selected 50 random samples of the data and marked channels that were correlated less than 85% to the other channels, were four standard deviations beyond the mean, had more line noise than signal, or were broken for more than half of the samples. Channels marked for rejection were then visually inspected in the raw data and then manually rejected if confirmed. These channels were interpolated using EEGLAB spherical interpolation. Sections of data were removed if they contained significant artifact or excessive movement. Data were then re-referenced to the common average before the second channel rejection step. During the second channel rejection step, two topographic plots were made for each dataset. One displayed the average amplitudes for the duration of the epoch and the other showed the difference amplitudes between conditions for the duration of the epoch. The spectrum of all channels was also plotted. These three plots were visually inspected for any additional bad channels (i.e. channels displaying exceedingly large amplitudes, surrounded by the opposite amplitude charge, or containing high/unusual spectral density). If any bad channels were identified, they were added to the list of bad channels for each dataset. Ultimately, the entirety of the first preprocessing stage resulted in a finalized list of channels to be rejected and sections of data to eliminate, both of which were used in the second and third stages of preprocessing.

During the second preprocessing stage, the raw data were prepared to be cleaned of artifacts via the weight transferring method ICA. First, flat-line channels were eliminated from the raw data and the data were filtered with a pass band of 1 to 45 Hz. Then the finalized list of bad channels was rejected (not interpolated), sections of data were eliminated, the data were re-referenced to the common average, and finally run through ICA to record the weights per channel. This second preprocessing stage resulted in a list of weights that would be transferred and assessed via “IClabel” in the third stage of preprocessing.

During the third preprocessing stage, flat-line channels were eliminated from the raw data, the data were filtered with a pass band of 0.1 to 45 Hz, the list of finalized bad channels were rejected (not interpolated), sections of bad data were eliminated, and then the data were re-referenced to the common average. ICA weights were transferred to these data, and components were rejected if they were labeled as less than 15% brain or more than 85% muscle, eye, heart rate, line noise, channel noise, or other artifact. Eliminated channels were then spherically interpolated. These final preprocessed data were used for further analysis. On average, these processing steps resulted in the removal of 14(±5) out of 128 channels, and 40(±24) out of 402 trials.

For analysis, epochs were generated from −200 to 1000 ms where baseline was the 200 ms before the onset of the final word in a sentence. Trials were automatically rejected if they breached the artifact rejection threshold of 250 μV or if the trial contained amplitudes larger than two standard deviations above the mean amplitude across channels. ERP waveforms were created by averaging across trials per condition at each electrode, and then across participants. Difference topography plots were calculated by subtracting the grand average amplitudes in response to congruent stimuli from the incongruent stimuli (incongruent – congruent amplitudes).

### Pre-planned analyses

JASP (Jeffrey’s Amazing Statistic Program Team [2020], Version 0.12.2) was used for statistical analyses. The primary outcome measures of interest here were the RP, N400 and the LPC/P600 because of their strong association with semantic processing, the construct of central interest in the current study. Prior research has identified the peak amplitude of the RP ∼250 ms over left frontal scalp regions in response to only congruent endings (best captured by measures at electrode F7) (Toffolo et al., 2022). The N400 ERP component has peak amplitudes at ∼400 ms after final word onset over midline scalp electrodes (typically represented by measures at Cz and Pz) (Colin Brown and Hagoort, 1993; Kutas and Hillyard, 1984; Osterhout and Holcomb, 1992; Toffolo et al., 2022). The subsequent response complex, the LPC/P600, typically peaks at ∼600 ms over left parietal scalp areas, best captured by measures at electrode P7 (Friederici, 2011; Kaan et al., 2000; Leckey and Federmeier, 2020; Toffolo et al., 2022; Van Herten et al., 2005). The grand average ERPs (+/- SEM) were therefore measured at four electrode sites (F7, Cz, Pz, and P7) during congruent and incongruent sentence presentation in both active (n=21) and passive (n=22) tasks (**Figure. 2**).

**Figure 1.**
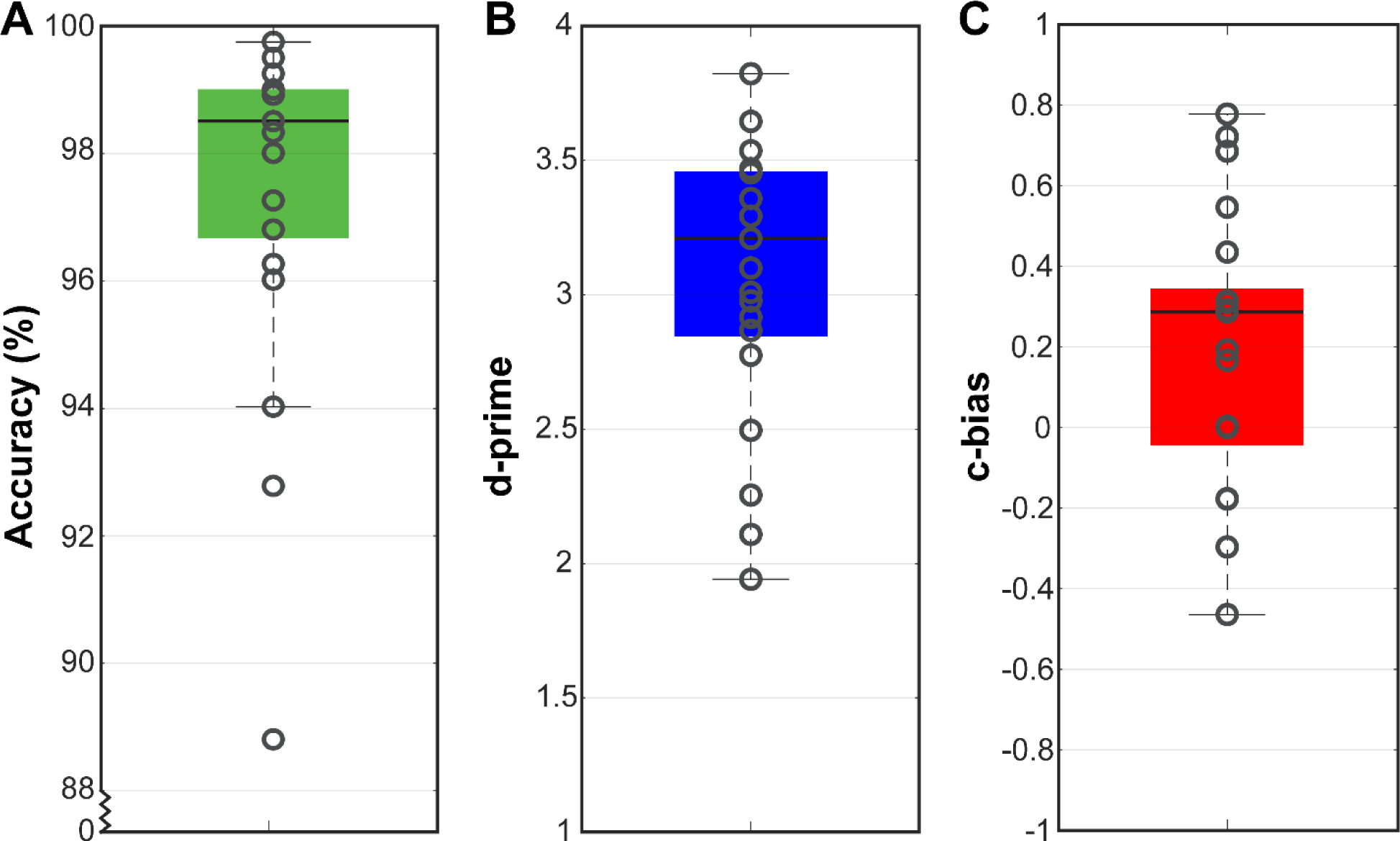
Accuracy and Behavior. Box and whisker plots showing the percent correct, d-prime, and c-bias of the subjects included in the active paradigm. The gray circles represent individual subjects, the black line is the median, the bottom and top edge of each box indicate the 25^th^ and 75^th^ percentiles respectively, and the extensions reach the most extreme subjects that are not considered outliers. Data points are considered outliers if they are beyond ±1.5 times the interquartile range plus the 75^th^ and 25^th^ percentile respectively. Positive c-bias values represent a bias towards subjects answering more incongruent trials correctly and congruent trials incorrectly.

**Figure 2.**
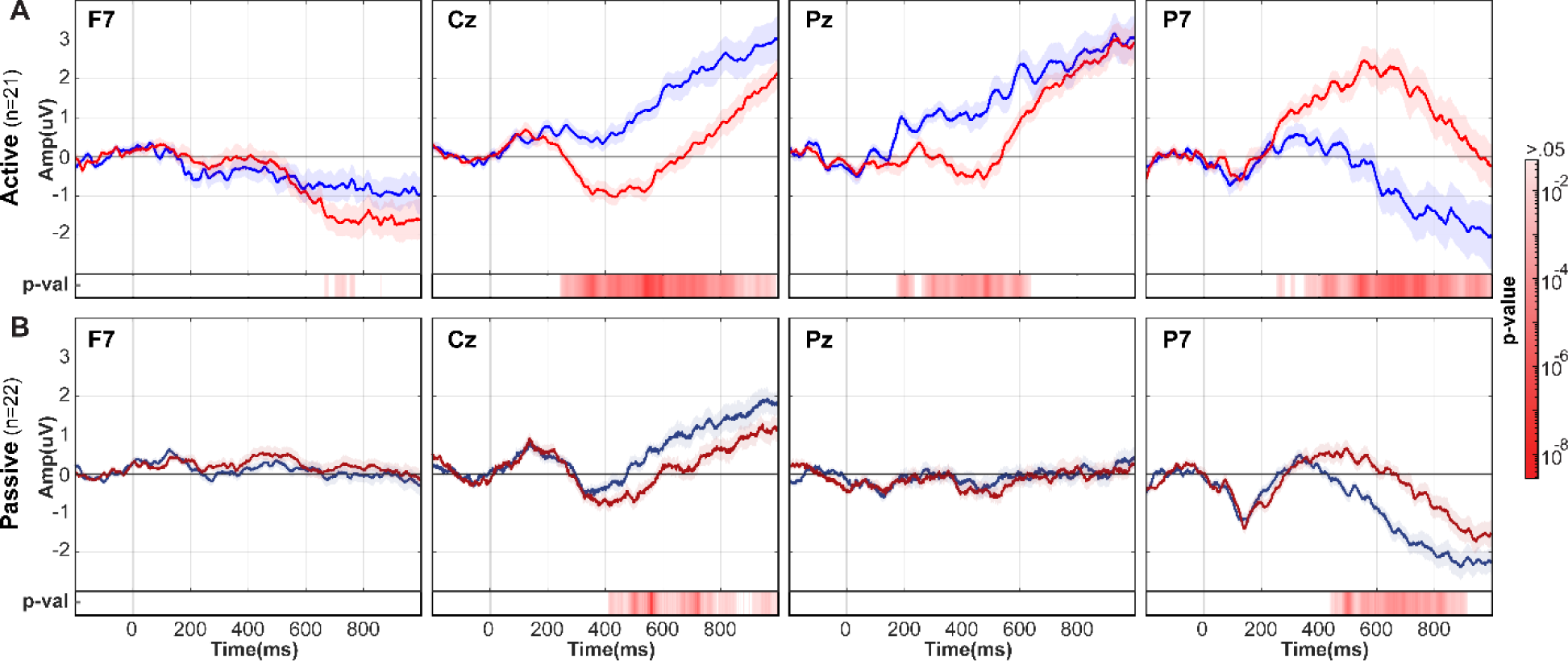
ERP Plots. ERP plots for frontal electrode F7, midline electrodes Cz and Pz, and left parietal electrode P7, show amplitude changes between −200-1000 ms for each condition (congruent vs. incongruent) and each paradigm (active vs. passive). Displayed are the waveforms in response to each condition during the **A.** active paradigm (blue, congruent; red, incongruent), **B.** and passive paradigm (navy, congruent; maroon, incongruent). The shading around the grand average amplitudes represents the standard error mean. Below each plot are p-values of the significant difference between conditions (white > 0.05) across the epoch derived from the spatial temporal cluster analysis.

Accordingly, *a-priori* analyses of the amplitude changes for the RP, N400, and LPC/P600 ERP components in response to different conditions (incongruent and congruent) during different paradigms (active and passive) were conducted at electrodes representing their expected scalp topographies and peak amplitude time points (i.e. electrode F7 at 250 ms for the RP, electrodes Cz and Pz at 400 ms for the N400 ERP, and electrode P7 at 600 ms for the LPC/P600) (Toffolo et al., 2022). Four separate 2 x 2 repeated measure ANOVA’s (rmANOVA) were conducted on the average amplitude values over 10 ms time-windows centered at 250 ms, 400 ms, and 600 ms, respectively (**Table 2**). Factors were named CONGRUENCE (congruent vs. incongruent endings) and ATTENTION (active vs. passive paradigms).

**Table 2.**
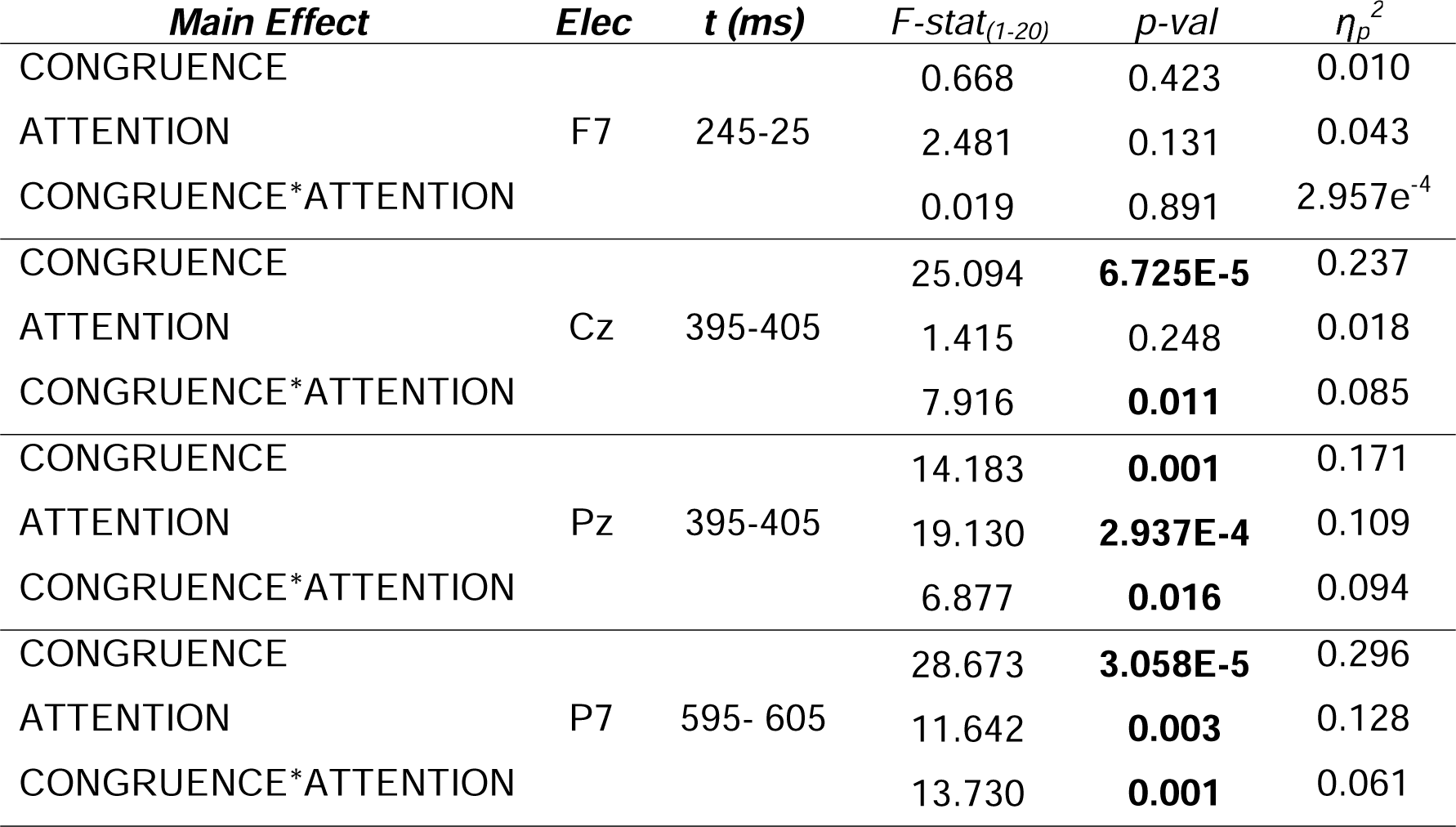
2 x 2 rmANOVA for frontal, midline, and left parietal electrodes. Amplitudes were acquired for both conditions during each paradigm by averaging the amplitude values from 5 ms before and after a time point of interest (i.e. 245-255, 395-405, and 595-605 ms). Using these values, the main effects of CONGRUENCE, ATTENTION, and CONGRUENCE by ATTENTION were assessed using a 2 x 2 repeated measure ANOVA (2 conditions by 2 paradigms) at each electrode separately (F7, Cz, Pz, P7). Bolded p-values represent a significant main effect (p<0.05, two tailed).

| *Main Effect* | *Elec* | *t (ms)* | *F-stat<sub>(1-20)</sub>* | *p-val* | $\eta_p^2$ |
| --- | --- | --- | --- | --- | --- |
| CONGRUENCE | F7 | 245-25 | 0.668 | 0.423 | 0.010 |
| ATTENTION |  |  | 2.481 | 0.131 | 0.043 |
| CONGRUENCE*ATTENTION |  |  | 0.019 | 0.891 | 2.957e<sup>-4</sup> |
| CONGRUENCE | Cz | 395-405 | 25.094 | **6.725E-5** | 0.237 |
| ATTENTION |  |  | 1.415 | 0.248 | 0.018 |
| CONGRUENCE*ATTENTION |  |  | 7.916 | **0.011** | 0.085 |
| CONGRUENCE | Pz | 395-405 | 14.183 | **0.001** | 0.171 |
| ATTENTION |  |  | 19.130 | **2.937E-4** | 0.109 |
| CONGRUENCE*ATTENTION |  |  | 6.877 | **0.016** | 0.094 |
| CONGRUENCE | P7 | 595- 605 | 28.673 | **3.058E-5** | 0.296 |
| ATTENTION |  |  | 11.642 | **0.003** | 0.128 |
| CONGRUENCE*ATTENTION |  |  | 13.730 | **0.001** | 0.061 |

### Exploratory Statistical Cluster Plot (SCP) analysis

In a second stage of exploratory analysis, the entirety of this rich spatio-temporal dataset was explored using the statistical cluster plot approach (Molholm et al., 2002). Describing the temporal course of the N400 and LPC/P600 components in this adolescent sample was a specific interest of this study because it was possible that the timing of these processes might not be fully mature in this cohort. Cluster plot statistics were generated using FieldTrip for MATLAB and displayed using the EEGLAB (Oostenveld et al., 2011). Group level cluster-based permutation tests were conducted using two-tailed, paired t-statistics with a critical alpha-level of 0.05 for each paradigm. The Monte-Carlo method was used to estimate significance probability along with triangulation spatial clustering and multiple-comparison corrections. Clusters were identified and combined using “maxsum” and a 5% two-sided cutoff criterion for both positive and negative clusters (**Figure 4**). To help visualize scalp areas that had significant differences between conditions, statistical topography plots were created by averaging the significance values in the cluster plots over a 10 ms time window centered at the time point of interest (**Figure 3D**).

**Figure 3.**
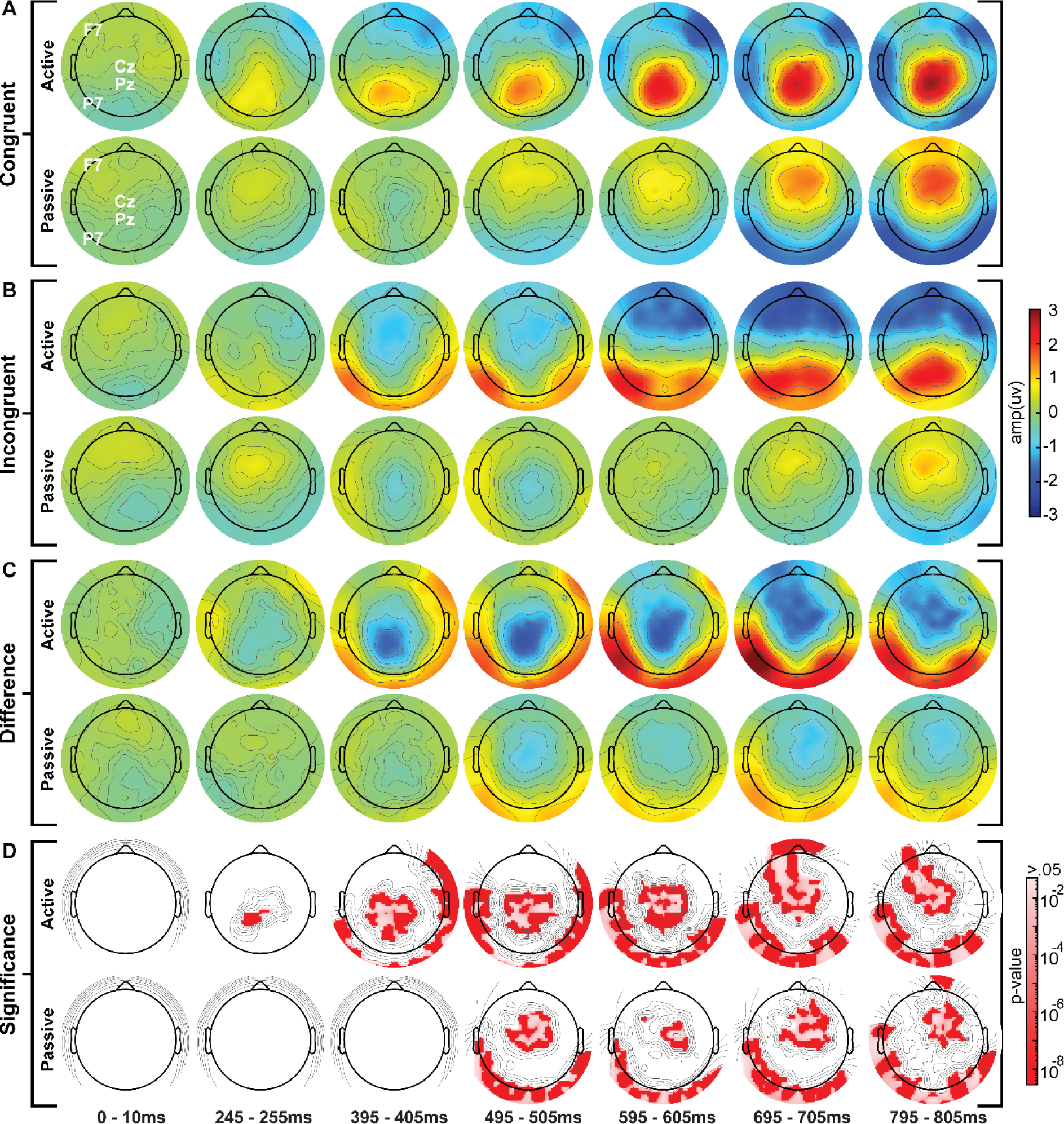
Grand average scalp topographies and p-values at 7 time points. The grand average distribution of EEG amplitudes for both active (n=21) and passive (n=22) paradigms averaged over 7 time windows (0-10 ms, 245-505 ms, 395-405 ms, 495-505 ms, 595-605 ms, 695-705 ms, and 8795-805 ms). Provided are the amplitudes for both paradigms (top, active; bottom, passive) in response to **A.** congruent stimuli and **B.** incongruent stimuli. Shown for both paradigms (top, active; bottom, passive) are **C.** the difference between conditions (incongruent-congruent amplitudes) and **D.** as a visual guide, the p-value cluster plots averaged over each time frame (red, p ≤ 0.05). Topographical electrode locations are shown on Figure 4A.

**Figure 4.**
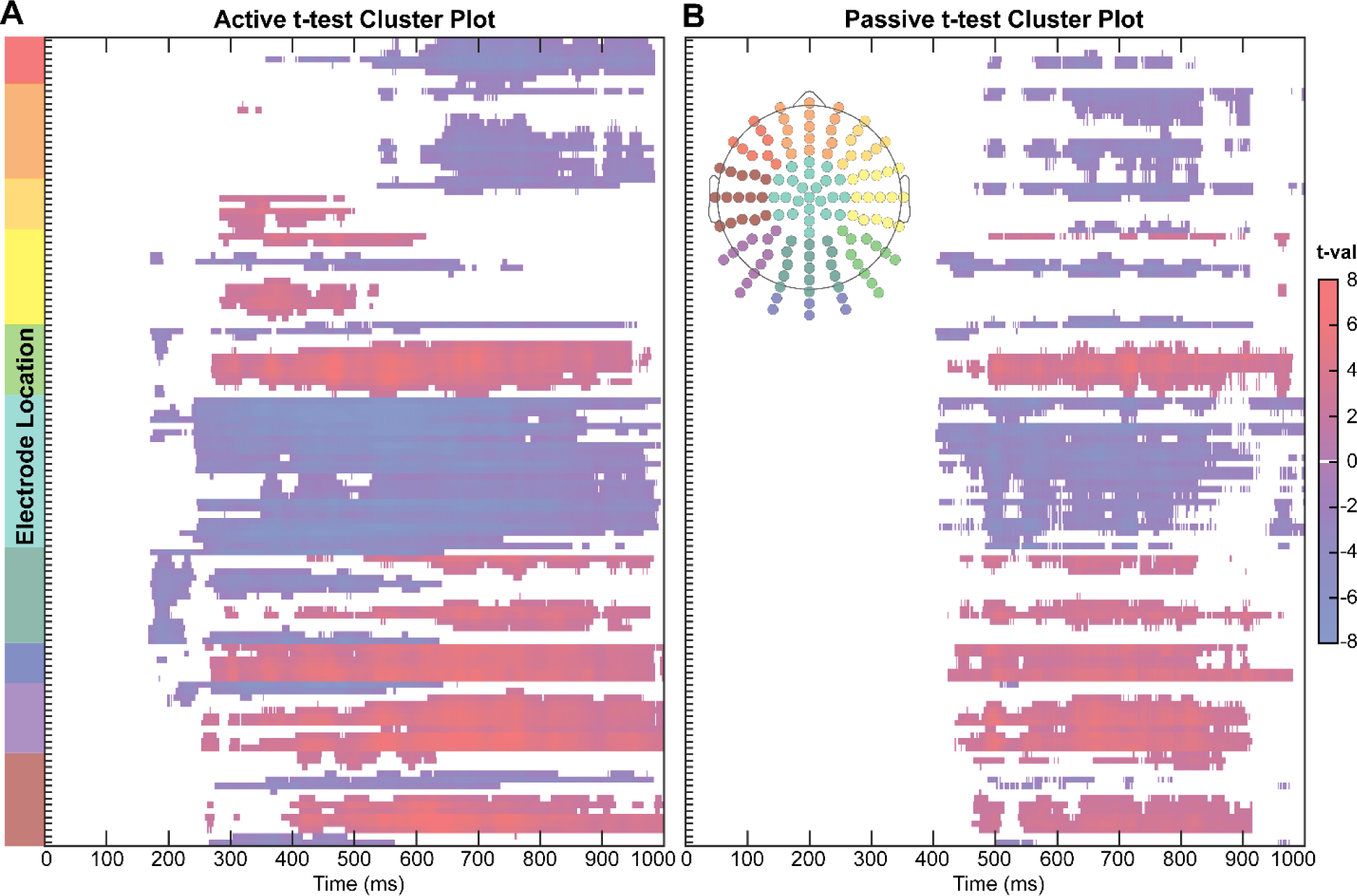
Spatial temporal cluster plots per paradigm. Cluster plots of two-tailed paired sample t-values of 128 channels over 1000 ms for **A.** active and **B.** passive paradigms. Cluster plots employ Monte-Carlo method t-tests with a 5% two-sided criterion cutoff, spatial clustering via the triangulation method, and a multiple-comparison correction. Electrode channels are arranged in the quadrants noted by the color coordinated scalp diagram.

### Post-hoc Topographic Dissimilarity and Microstate Analyses

A *post-hoc* analysis was conducted to assess for possible significant topographic differences between the active and passive responses to each condition. This was achieved by first calculating the normalized global field power (GFP) for each condition (congruent or incongruent) and paradigm (active or passive). The GFP for each time point was acquired by taking the square root of the sum of the squared mean amplitudes across all electrodes, divided by the total number of electrodes (Skrandies, 1990). Topographical differences were then quantified via global dissimilarity (DISS) at the individual participant level. DISS is an algorithm to calculate the distance between two GFP-normalized vectors (i.e. electric field topographies) at a given time point. DISS was calculated between the responses to each condition (i.e. active congruent vs. passive congruent, or active incongruent vs. passive incongruent). Specifically, at each time point, the square root of the mean of the squared differences between potentials was measured at each electrode and divided by the GFP. The DISS index for a time point ranges between 0, identical topographic distribution, and 2, reversed topographic distribution. The significance of these topographic dissimilarities was then assessed via a non-parametric bootstrapping procedure, the so-called topographical ANOVA (TANOVA). First, the mean DISS and standard deviation for the null hypothesis (i.e. no differences between the topographies of a given pair of conditions (e.g. active congruent versus passive congruent, and all other pairings of conditions)) was acquired through 2000 permutations of shuffling subjects between paradigms (active and passive) of the same condition (e.g. congruent) and calculating the DISS scores between the resulting randomized groups (Lehmann and Skrandies, 1980; Murray et al., 2008; Skrandies, 1993). Second, the z-statistic was calculated per time point of a single condition by taking the observed DISS index, subtracting the mean null DISS, and dividing by the standard deviation of the null DISS. A clustering procedure was then applied to only include significant time points with a p-value ≤ 0.05 (z < −1.645 and z > 1.645) and if that significant time point was part of a cluster that remained significant for at least 11 consecutive time points.

To further characterize the observed topographical differences, a microstate analysis was conducted on the data using the Microstate EEGLAB toolbox (Poulsen et al., 2018). Using this toolbox and its step-by-step instructions, the ERPs of all subjects regardless of condition or paradigm were pooled to construct microstate prototypes. The selected statistical cluster analysis was the Topographic Atomize and Agglomerate Hierarchical Clustering (TAAHC), which prioritizes topographic similarity (global explained variance, GEV) over field strength (GFP) (Poulsen et al., 2018). Data were normalized before segmentation and polarity was taken into account. Four clusters were selected based on the displayed GEV and cross-validation criterion (CV) scores, which were high and low respectively (i.e. adding more clusters would not benefit the results). The four prototype microstates were then back-fit onto the grand average ERP data for each condition and paradigm, and segmented across their respective GFPs. Segments were rejected if the duration of the microstate was less than 30 ms. The average duration and GEV a microstate had relative to each condition and paradigm were acquired via the “pop_micro_stats” function.

### Data Availability

The stimuli and supporting datasets for this experiment can be freely accessed on OpenNeuro: ds004940 (https://openneuro.org/datasets/ds004940). The dataset download provides 22 datasets organized in BIDS format via guidelines provided by (Pernet et al., 2019), participant information (.tsv and .json), and stimulus set files. Included in the stimulus set are the auditory files for all 404 stimuli and the auditory files for introductory explanations and practice trials. Additionally, a stimulus parameter file is provided, which includes stimulus information such as duration, target word onset, and the written form of each stimulus so that reading semantic comprehension can also be investigated.

The full dataset includes unfiltered EEG data (.bdf) for both paradigms (active and passive), corresponding event files, and channel rejection files for each participant (.tsv), as well as recording information, electrode positioning, and event file information (.tsv and/or .json). Derivatives are also provided for transparency. Derivatives include files that were used for manual filtering decisions (.mat, .fig, and .tsv), preprocessed EEG data filtered with ICA (.mat), and preprocessed grand average ERP data for this study (.mat) along with the corresponding trial rejection information for each participant (.tsv), and filtering parameters (.json). Refer to the README file in order to use the dataset appropriately. Use of this dataset, stimulus set, or presenting examples from this stimulus set are conditioned upon appropriate attribution (i.e. citation of this paper).

The original stimulus set (https://doi.org/10.5061/dryad.9ghx3ffkg), which includes all 442 stimuli and the cloze probability survey from our previous study (Toffolo et al., 2022), along with the original 24 adult observer datasets (https://doi.org/10.5061/dryad.6wwpzgmx4) are available through Dryad for the scientific community to use freely.

### Code Availability

The code generated for the analysis of these datasets can be freely accessed on OpenNeuro: ds004940 (https://openneuro.org/datasets/ds004940). The dataset download provides code that was used to preprocess the EEG data, generate ERPs, and create components of each figure. The Presentation® code used in the experiment is also provided. The analysis code is only usable on datasets in BIDS format. Please refer to the README in order to use this code appropriately.

## RESULTS

### Behavioral Performance

During the active response task, the 21 adolescents included in these analyses correctly identified congruent and incongruent sentence endings with 97.5% accuracy (± 2.7%). **Figure 1** illustrates the distributions of response accuracy (**Figure 1A**), d-prime (sensitivity index) (**Figure 1B**) and C-bias (decision criterion) (**Figure 1C**). During the passive presentation task, participants did not indicate whether they thought sentences were either congruent or incongruent.

### ERP Components in Active and Passive Contexts

Displayed in Figure 2 are the amplitude values during the active (Figure 2A) and passive (Figure 2B) paradigm in response to each condition (Congruent: Red-Active, Maroon-Passive; Incongruent: Blue-Active, Navy-Passive) over left frontal, midline, and left parietal scalp, for which time point 0 represents the onset of the last word of a sentence. Below each plot are bars of significance for each electrode, derived from a *post-hoc* spatial-temporal cluster plot analysis. Here, red represents significant *p*-values while white indicates *p*-values > .05.

### The Recognition Potential (RP)

A 2 x 2 rmANOVA with factors of CONGRUENCE and ATTENTION (i.e. congruent vs. incongruent, and active vs. passive) was conducted on mean amplitude measures taken from the left frontal electrode site (F7) for the 245-255 ms time-window. The outcomes of this analysis are reported in full in **Table 2**. There were no main effects of CONGRUENCE, ATTENTION, and no interaction of CONGRUENCE by ATTENTION, suggesting that these semantic stimuli did not evoke the RP in this sample of adolescents (all *p-*values >0.13). Observationally, over left frontal scalp, regardless of condition, there was a slow negative-going potential beginning around stimulus onset during the active paradigm, and minimal change in amplitude during the passive paradigm (**Figure 2**).

### The N400 component

Two 2 x 2 rmANOVA’s were conducted on mean amplitude measures taken from the 395-405 time window at the midline electrodes Cz and Pz. Again, full results of this analysis are shown in **Table 2**. A main effect of CONGRUENCE (Cz: F_(1-20)_=25.094, *p*=6.725e^-5^, η_p_^2^=0.237; Pz: F_(1-20)_=14.183, *p*=0.001, η_p_^2^=0.171) and an interaction of the factors of CONGRUENCE and ATTENTION (Cz: F_(1-20)_=7.916, *p*=0.011, η_p_^2^=0.085; Pz: F_(1-20)_=6.877, *p*=0.016, η_p_^2^=0.094) wereobserved. There was also a main effect of ATTENTION at Pz (F_(1-20)_=19.130, *p*=2.937e^-4^, η_p_^2^=0.109). The effect of CONGRUENCE was reflected in the ERP response waveforms over central and parietal scalp (**Figure 2**). Here, in response to incongruent endings relative to congruent endings, there was a significantly larger negative deflection between 200 and 600 ms, with a distinct peak ∼400 ms regardless of paradigm (i.e. the N400). A significant difference between conditions was not just present at this timeframe, but was maintained through the entirety of the epoch. Over parietal scalp, the effect of ATTENTION alone was visible. Amplitudes were more positive during the active paradigm relative to the passive paradigm, for which there was little change in response amplitude regardless of condition. An interaction of CONGRUENCE by ATTENTION was also observed over central and parietal midline scalp in that amplitude differences between conditions (i.e. the N400 effect) around 400 ms were larger in the active paradigm relative to the passive paradigm over both central and parietal scalp (**Figure 2A** and **2B**). Furthermore, amplitudes in response to congruent endings were more positive at this time during the active paradigm (**S1**).

### The Late Positivity Component (LPC/P600)

A 2 x 2 rmANOVA with factors of CONGRUENCE and ATTENTION was conducted on amplitude measures taken from left parietal electrode P7 at 595-605 ms. Here, there were significant main effects of CONGRUENCE (F_(1-20)_=28.673, *p*=3.058e^-5^, η_p_^2^=0.296), ATTENTION (F_(1-20)_=11.642, *p*=0.003, η_p_^2^=0.128), and CONGRUENCE by ATTENTION (F_(1-20)_=13.730, *p*=0.001, η_p_^2^=0.061). Full results of this analysis are also shown in **Table 2**. The effect of CONGRUENCE was observed in the ERP waveforms ∼600 ms over left parietal scalp. Here, in response to incongruent endings relative to congruent endings, there was a significantly larger positive deflection between 200-1000 ms that peaked ∼600 ms regardless of paradigm (**Figure 2**). The effect of ATTENTION was also visible in that regardless of condition, amplitudes during the active paradigm were more positive than the passive paradigm in this time frame. Lastly, the interaction effect of CONGRUENCE by ATTENTION was also observed. Amplitude differences between conditions (i.e. LPC/P600 effect) were greater in the active paradigm than the passive paradigm (**Figure 2A** and **2B**), specifically, the response to incongruent endings was more positive ∼600 ms in the active paradigm (**S1**).

### Scalp Topographic Mapping

The group averaged topographies for each paradigm are illustrated during seven 10 ms epochs (0-10 ms, 245-255 ms, 395-405 ms, 495-505 ms, 595-605 ms, 695-705 ms, and 795-805 ms) to show the evolution of the various processing stages across time (**Figure 3**). Here, red signifies more positive amplitudes and blue more negative amplitudes.

### Topographic distributions in the Recognition Potential (RP) timeframe

During the active paradigm between 245-255 ms, there were slight negative amplitudes (∼1μv) over frontal scalp in response to congruent stimuli (**Figure 3A**), and little amplitude changes over frontal scalp in response to incongruent stimuli (**Figure 3B**). As a result, there was a small positive amplitude difference between these conditions over right frontal scalp at this timeframe (245-255 ms) that was non-significant (**Figure 3C** and **D**). Alternatively, during the passive paradigm at the same time frame, there were little changes in amplitude across the scalp in response to congruent endings, and minor fronto central positivity (∼1μv) in response to incongruent endings. There were no differences between conditions at this time frame during the passive paradigm.

### Topographic distributions in the N400 timeframe

In response to congruent sentence endings, during the active paradigm, positive amplitudes (between 2-3 μV) were present over central parietal scalp between 395-405 ms and subsequent time points. During the passive paradigm, positive amplitudes were not only seen at later time points than the active paradigm (starting between 595-605 ms and later), but also over more fronto-central scalp, quite distinct from the central-parietal positivity during the active paradigm (**Figure 3A**).

In response to incongruent endings during the active paradigm, negative amplitudes were broadly present over central scalp locations between 395-405 ms and 495-505 ms, highly consistent with where the N400 effect is typically observed (C. Brown and Hagoort, 1993; Kutas and Hillyard, 1984; Osterhout and Holcomb, 1992; Toffolo et al., 2022). This negativity shifted to frontal scalp areas at later time points. During the passive paradigm, negative amplitudes were seen over central scalp, although attenuated relative to the active paradigm and only present between the 395-405 ms and 495-505 ms time points.

Topographical difference plots (incongruent - congruent amplitudes) and significance plots for each paradigm are shown in **Figure 3C** and **D**, respectively. During the active paradigm, there was a significant negative difference between conditions between 395-405 ms (i.e. N400 effect) over central scalp. This significant central negative difference was maintained in all subsequent time frames, with more central-parietal scalp distribution early in the epoch (between 245-605 ms) and fronto-central distribution later in the epoch (between 695-805 ms). Relative to the active paradigm, the passive paradigm had strikingly lower amplitude differences across the scalp. A significant negative difference over central parietal areas was not present between 395-405 ms during the passive paradigm as it was during the active paradigm. Alternatively, a more localized significant fronto-central negative difference was seen between 495-505 ms and at later time periods.

### Topographic distributions in the Late Positivity Component (LPC/P600) timeframe

In response to congruent sentence endings, during the active paradigm, there were positive amplitudes (circa 2-3 μV) present over central-parietal scalp between 595-605 ms and subsequent time points. There were also negative amplitudes over right frontal scalp (circa −1.5 μV) between 595-605 ms, and negative amplitudes over posterior and left temporal locations at later time points. During the passive paradigm, the positive amplitudes (∼1μv) were instead over fronto-central scalp during this time frame (595-605 ms) and subsequent time points. Frontal negativity was not observed at this time, but there were negative amplitudes over posterior locations between 695-705 ms and 795-805 ms (**Figure 3A**).

In response to incongruent sentence endings, during the active paradigm, positive amplitudes were present over temporo-parietal and posterior scalp starting between 395-405 ms, with slightly larger amplitudes over the left hemisphere. Positive amplitudes were more robust at later time points over temporo-parietal and posterior scalp, where the LPC/P600 ERP is commonly observed (Friederici, 2011; Kaan et al., 2000; Leckey and Federmeier, 2020; Toffolo et al., 2022; Van Herten et al., 2005). During the passive paradigm, positive amplitudes were not present over posterior scalp areas at any time point. Unlike the active paradigm, at later time points during the passive paradigm, there were fronto-central positive amplitudes and posterior negative amplitudes (**Figure 3B**). The conditional responses during the passive paradigm were very similar between 795-805 ms, although with reduced amplitudes in response to the incongruent condition relative to the congruent condition (**Figure 3A, Figure 3B**).

During the active paradigm, there were significantly positive differences between conditions over lateral and posterior scalp (**Figure 3C** and **D**). Starting between 395-405 ms, this significant positivity was lateralized to the right hemisphere, but was more posterior and bilateral at later time points. During the passive paradigm, this significant positive lateral difference between conditions appeared later than the active paradigm (between 495-505 ms), with a more bilateral posterior scalp distribution, and with reduced amplitudes (**Figure 3C** and **D**).

### Exploratory Spatial-Temporal Cluster Plots

Spatial-temporal statistical cluster plots for each paradigm across the entire epoch are shown in **Figure 4**. Red represents significantly positive t-values, blue represents significantly negative t-values, and white indicates non-significant t-values. These t-test cluster plots recapitulate what was seen in the *p*-value topographic scalp maps (**Figure 3D**), but t-values are provided for every time point across all electrodes during the epoch. Overall, significant differences between conditions were found over less time points throughout the epoch during the passive paradigm relative to the active paradigm. During the active paradigm (**Figure 4A**), there was a significant negative difference over central-parietal scalp starting as early as 200 ms after target word onset, that was maintained for nearly the entirely of the epoch. Alternatively, during the passive paradigm (**Figure 4B**), a significant negative difference between conditions started later (i.e. ∼450 ms), was not sustained as long as the active paradigm (ending ∼900 ms), and was primarily localized over frontal-central scalp. A significant positive difference was present during both paradigms over bilateral parietal and posterior scalp. However, this positivity began ∼150 ms earlier in the active paradigm (∼300 ms) relative to the passive paradigm and was sustained for the remainder of the epoch. There was also significant positive activity over right frontal electrode sites between 300 and 500 ms during the active paradigm.

### Topographic Dissimilarity Analysis

The topographical differences between the active versus passive response to a single condition are shown in **Figure 5**. The top panels show the GFP for each paradigm in response to the congruent condition (**Figure 5A**) and incongruent condition (**Figure 5B**), while the bottom panels show the significant DISS scores between the paradigms across the epoch. Significant differences between the topographical spatial orientations via a TANOVA were found primarily between ∼100-250 ms and ∼300-1000 ms in response to the congruent condition (**Figure 5A**), and between ∼150-350 ms and ∼400-1000 ms in response to the incongruent condition (**Figure 5B**). Further analysis of this topographical difference was explored via a microstate analysis (**Figure 6**). Active and passive microstate segmentation were different from each other. During the active paradigm, there was an early frontal positivity and posterior negativity seen in the topographic responses to both conditions (microstate 2). However, ∼200 ms, this transitioned into a microstate dominated by central scalp positivity and frontal negativity in response to the congruent condition (microstate 3), while in response to the incongruent condition, the data were most reflective of microstate 4, with fronto-central negativity and left temporal-occipital positivity (**Figure 6A** and **B**). Unlike the active paradigm, the microstates and their transitions were very similar between the conditions of the passive paradigm (**Figure 6A**). Like the active paradigm, early periods of these passive epochs were defined by microstate 2. However, this microstate was present for longer between 0-300 ms. This earlier period was also defined by larger GFPs than the active paradigm, which peaked around 150 ms. For both conditions of the passive paradigm, albeit slightly longer for the incongruent condition, microstate 2 transitioned briefly into microstate 4 before returning to microstate 2 (**Figure 6A** and **B**). Both conditions in the passive paradigm were in microstate 4 at 400 ms like the active incongruent condition. Values for the average time spent in and similarity to each microstate for each condition and paradigm are shown in **Figure 6C**. The topography over the active congruent epoch was most similar to microstate 1 and 3, with positive amplitudes over broad central scalp regions, but negative amplitudes over frontal and bilateral temporal-occipital electrodes respectively. Alternatively, the topography over the active incongruent epoch was most similar to microstate 3 and 4. The topography during both conditions of the passive paradigm were highly similar to microstate 2, with positivity over frontal scalp electrodes and negative amplitudes over posterior scalp.

**Figure 5.**
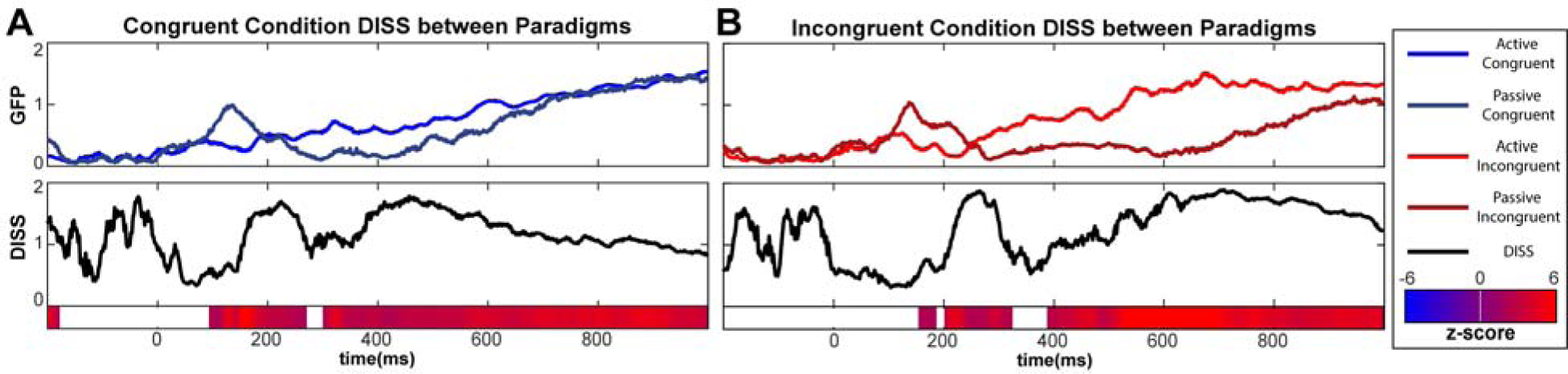
Topographical Differences using Global Dissimilarity Index. Topographical differences between paradigms (active vs. passive) in response to each condition (congruent and incongruent). The top panels show GFP over 1200 ms in response to **A.** the congruent condition (blue, active; navy, passive) and **B.** the incongruent condition (red, active; maroon, passive). The 0 ms marks the onset of the last word in a sentence and the 200 ms prior is baseline. The bottom panels show the raw DISS in black, with corresponding z-scores below for the whole epoch. Blue represents more negative z-scores and red more positive z-scores. White represents non-significant z-scores (> −1.645 and < 1.645) for the DISS between the paradigms of a single condition.

**Figure 6.**
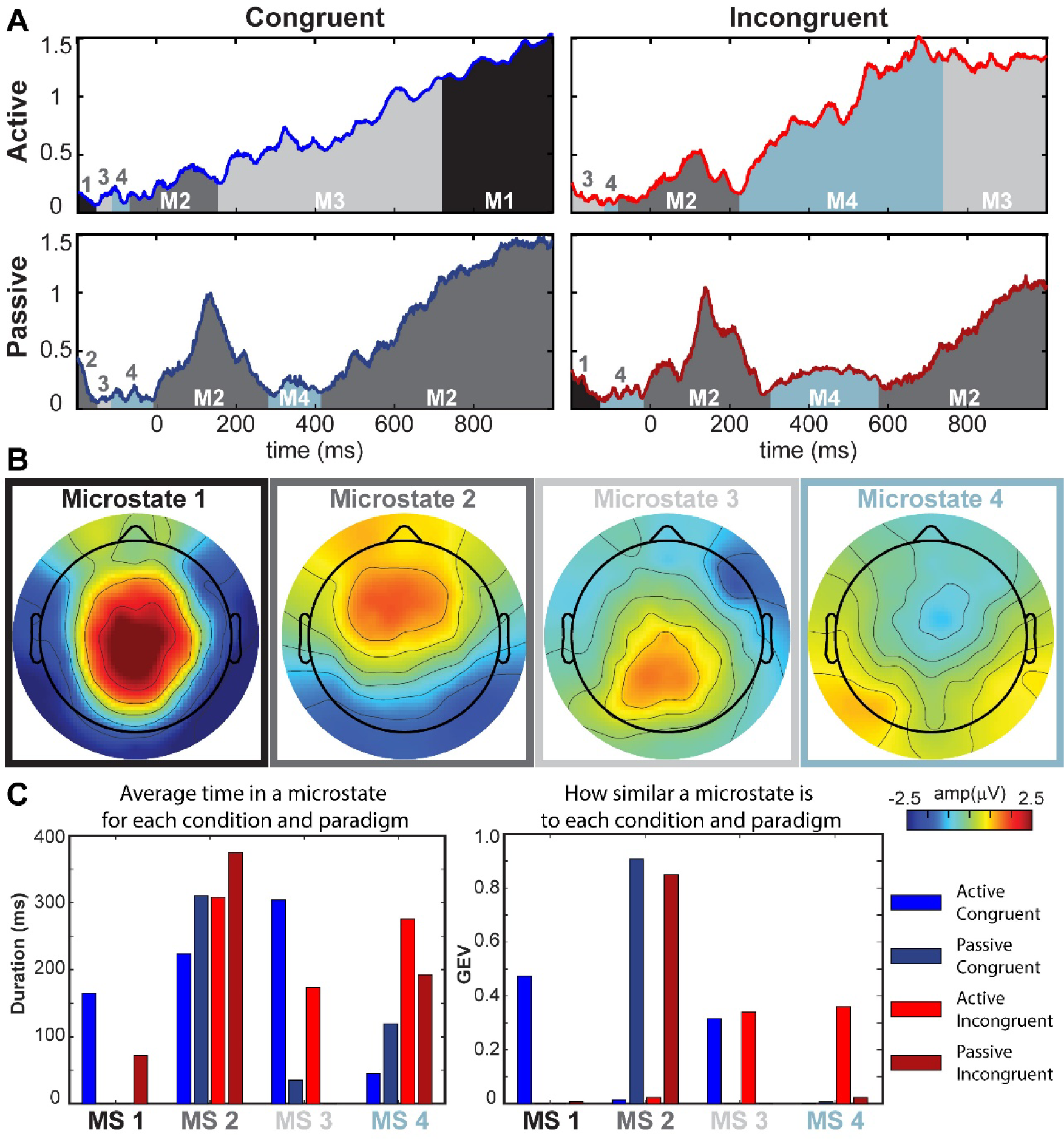
Microstate Analysis. Microstate analyses of the ERPs evoked in response to congruent and incongruent conditions, during active and passive paradigms. **A.** GFP plots over 1200 ms for each condition and paradigm. The color shading beneath each GFP corresponds to the time periods for which the topography of a condition and paradigm was most similar to one of the 4 microstates. **B.** Displays the 4 prototypical microstates derived from the concatenation of all subject ERPs for each condition and paradigm after conducting a topographic pattern analysis. Red signifies more positive potentials and blue more negative potentials. **C.** Shows the total duration that a microstate had for each condition and paradigm (left), and how similar each microstate was to each condition and paradigm through GEV scores (right).

## Discussion

We set out here to investigate whether higher-level sentential semantic comprehension could be measured in passive listening contexts, with an eye to developing a tool for use in pediatric populations with communication differences, such as disorders of consciousness (DOC), minimally verbal autism spectrum disorder (mvASD) and Rett-Syndrome. Consistent with prior literature and our main hypotheses, attention affected the processing of semantic congruency, substantially modulating the amplitudes of the N400, as well as the LPC/P600 in a sample of 22 neurotypical adolescents between the ages of 12 and 17 years.

When actively attending the stimuli, there were significantly larger amplitude deflections around 400 ms over central-parietal regions of the scalp in response to incongruent sentence endings relative to congruent sentence endings, indicative of the N400 ERP component (Block and Baldwin, 2010; Kutas and Hillyard, 1984, 1980; Lau et al., 2008; Luck, 2005; Osterhout and Holcomb, 1992; Toffolo et al., 2022). Semantically driven responses were also seen at later time points over posterior scalp. In response to incongruent sentence endings, there was a significantly larger positive amplitude deflection that peaked at 600 ms, indicative of the LPC/P600. However, comparing across studies, there were observable differences in the adolescent N400 relative to adults who experienced the same stimuli in our prior study (Toffolo et al., 2022). The reader is reminded that this earlier study with adults employed exactly the same paradigm and recording parameters as the current adolescent study. The amplitudes of the N400 and LPC/P600 in adolescents were twice that of adults, and the N400 effect (i.e., significant differences between conditions ∼400 ms) over central scalp showed a more protracted latency in adolescents. Here, rather than a N400 effect between 200-600 ms exhibited by adults, the N400 effect in adolescents started ∼200 ms and was extended to 1000 ms after the onset of the final word in a sentence. Furthermore, the RP, which is thought to represent the point at which an individual recognizes the target word following congruent context, was not detected over left frontal scalp in this adolescent sample, suggesting that context prediction abilities may not be fully developed in adolescents (Fernandez and Smith Cairns, 2011). The very high accuracy levels observed behaviorally during the active paradigm showed that neurotypical adolescents fully understood the meaning of the sentences, so differences in amplitude, latency, and presence of these ERP components relative to adults are likely a function of still maturing adolescent language processing rather than differences in semantic processing abilities. Regardless of these differences, clear N400 and LPC/P600 ERP components were evoked in neurotypical adolescents during the active paradigm.

Similar waveforms were elicited when ignoring the stimuli, but there was a significant interaction between the factors of CONGRUENCE and ATTENTION over central and posterior midline scalp sites, pointing to clear influences of attention on these semantic processing ERP components. Relative to the active paradigm, the ERPs evoked during the passive paradigm were attenuated in amplitude, and significant differences between conditions were present over less extensive areas of scalp, both of which resulted in a delayed onset of conditional differences (i.e. N400 and LPC/P600 effects) in the passive paradigm. The N400 ERP component was clearly elicited over central midline scalp in the passive paradigm, but not posterior midline scalp. Additionally, there was a larger amplitude response to congruent stimuli at 400 ms, which resulted in a reduced amplitude and delayed N400 effect. The amplitudes of the LPC/P600 in response to incongruent stimuli were also attenuated and delayed over left posterior scalp areas.

In addition to these clear differences in N400 and LPC/P600 component amplitudes and durations as a function of attention, the topographic distributions in response to both conditions were also found to be quite distinct, pointing to the fact that the underlying circuitry engaged for processing semantic information was fundamentally different under active versus passive contexts. For example, in response to actively attended congruent stimuli, a prominent midline central-parietal positivity with a concomitant right lateralized frontal negativity were observed during the N400 timeframe. In contrast, when the same congruent stimuli were heard under passive conditions, there was a later onset of fronto-central positivity with bilateral occipito-temporal negativities (see panel A of Figure 3). Fully distinct topographic distributions were also evident for incongruent stimuli under active versus passive conditions. Post-hoc topographic statistical analyses confirmed what was visually evident from the maps - that these distributions were indeed statistically distinct. In our initial hypotheses, we did not predict differential generator configurations under the different attention conditions, so this post-hoc observation will need to be followed up and confirmed in future work before any strong conclusions can be derived. Nonetheless, these findings suggest that the fundamental processing of semantic information is different under passive versus active contexts.

While withdrawal of explicit attention from the sentence stimuli resulted in attenuation of the N400 and LPC/P600 effects in the current data, these components were still robustly evoked in this adolescent cohort under passive listening conditions. Thus, in a setting where semantic inputs were essentially being ignored in favor of a simultaneous TV show, semantic congruence was nonetheless being monitored in the “background”. It is important to acknowledge that the extent of engagement with the visual TV input was not objectively measured here, so it is possible that attention could have drifted to the auditory inputs on occasion, or that attention was partially divided between sensory streams. A stronger manipulation would have been to require participants engage in an active orthogonal visual task that was psychophysically titrated to require strong attentional engagement to perform. Such a paradigm might well “force” a suppression of auditory processing in favor of resource allocation to the visual task (Foxe et al., 2005; Foxe and Simpson, 2005), and might have further reduced or even eliminated these semantic processing ERP components. However, our purpose here was tool development, and not to determine if covert semantic processing could be eliminated through intersensory selective attention mechanisms. As pointed out above, it is of significant value to be able to objectively measure covert semantic processing abilities in clinical populations where communication is difficult or impossible, and explicit task performance cannot be obtained. In our experience, a highly feasible way to acquire high-density EEG from individuals with intellectual or neurodevelopmental disorders is to have them watch a favorite TV show during collection (Brima et al., 2024b; Francisco et al., 2020; Isenstein et al., 2024). As such, it is a reasonable proposition that mimicking this scenario in neurotypical adolescents can provide valuable insights into whether a passive experience of background sentential inputs can induce key neural markers of semantic processing (i.e. N400 and LPC/P600), which is clearly the case here.

### Study Limitations

As with any study, there are limitations that should be pointed out. Data from only 22 participants across a relatively wide age-range (12-17 years) were used for this study. A larger dataset would be required to adequately model the impact of development across this timeframe, and data from younger children would be ideal for mapping the developmental trajectory of the processes of interest. Additionally, this study presents each sentence twice to a participant, once with a congruent ending and once with an incongruent ending. While the order of congruence versus incongruence for any give sentence is randomized, this does introduce the possibility of repetition effects. This was directly examined in our previous work in adults using the same stimulus set, and it was found that order of presentation did not detectably affect the ERP components of interest (Toffolo et al., 2022). The stimuli in this set were also non-prosodic (i.e., monotone). This is important for studying the semantic comprehension of different populations, especially populations known to have difficulty with prosody, such as ASD or Schizophrenia (DePape et al., 2012; Leitman et al., 2007, 2005; Martzoukou et al., 2017; McCann et al., 2007), because it ensures that controls will not have an advantage in language comprehension. However, the lack of prosody may pose a limitation for this study, especially during the passive paradigm. Here, non-prosodic sentences may reduce the level of automatic engagement as opposed to prosodic sentences. Despite the potential for reduced automatic engagement with these stimuli, there still was a significant difference between conditions during the passive paradigm. With this in mind, it is possible that the presentation of prosodic sentences in a passive context will result in larger N400 and LPC/P600 effects, something to be explicitly explored in future work. Another consideration is that we limited the sample to individuals who reported English as their first language. As such, the current results should not be generalized to individuals whose primary language is English, but who had acquired a different language first. Otherwise, this study sought to have participants reflective of the general population, including both left-handed and bi- or multi-lingual individuals. Participants in both of these groups exhibited N400 and LPC/P600 ERP components (**S2**).

## Conclusions

This study successfully elicited neurophysiological indices of semantic processing in adolescents aged 12-17 years. When actively attending sentences with semantic errors, the N400 ERP component and the LPC/P600 were evoked, but not the RP. And although attenuated and with different spatial topography, there were significant differences in the amplitudes of these ERPs in response to sentences with and without semantic errors, even when attention was diverted away from the stimuli. These data suggest that the stimulus set used in this study could be used to objectively investigate semantic processing in populations with communication deficits.

## DECLARATIONS

### Ethics and Consent

The Research Subjects Review Board of the University of Rochester approved all the experimental procedures (STUDY00002036). Each participant provided written informed consent in accordance with the tenets laid out in the Declaration of Helsinki.

## Acknowledgments

The authors acknowledge the support of the autism team at the Frederick J. and Marion A. Schindler Cognitive Neurophysiology Lab, which includes Erin Bojanek, Emily Isenstein, Emily Knight, Paige Nicklas, and Zachary Christensen. We are especially grateful to the participants in this study who graciously gave their time to allow for the successful completion of this work.

## Authors’ Contributions

All authors collectively conceived of the study. KKT recruited participants and collected the data. KKT analyzed the data in consultation with EGF and JJF, and produced the figures for this study. KKT wrote the first draft of the manuscript and all authors provided editorial input on subsequent drafts. All authors read and approved the final version of this manuscript.

## Data Sharing

The stimuli, supporting datasets, and code for this experiment can be freely accessed on OpenNeuro: ds004940. The dataset download provides 22 datasets organized in BIDS format via guidelines provided by (Pernet et al., 2019), participant information (.tsv and .json), and stimulus set files.

## Consent for publication

Not applicable

## Abbreviations List

ASD: Autism Spectrum Disorder
mvASD: Minimally Verbal Autism Spectrum Disorder
CMS: Common Mode Sense
DISS: Global Dissimilarity Index
DOC: Disorders of Consciousness
EEG: Electroencephalography
ERP: Event Related Potential
GFP: Global Field Power
GEV: Global Explained Variance
ICA: Independent Component Analysis
LPC: Late Positivity Component
NT: Neurotypical
RP: Recognition Potential
SEM: Standard Error Mean

