## Supplemental Files for "Neurophysiological measures of covert semantic processing in neurotypical adolescents actively ignoring spoken sentence inputs: A high-density event-related potential (ERP) study"

| **Table of Contents:** | **Description** | **Pages** |
| --- | --- | --- |
| Figure S1. | ERP Plots by Condition | 1 |
| Figure S2. | Individual Subject Responses in the Active Paradigm | 2-7 |
| Figure S3. | Individual Subject Responses in the Passive Paradigm | 8-12 |

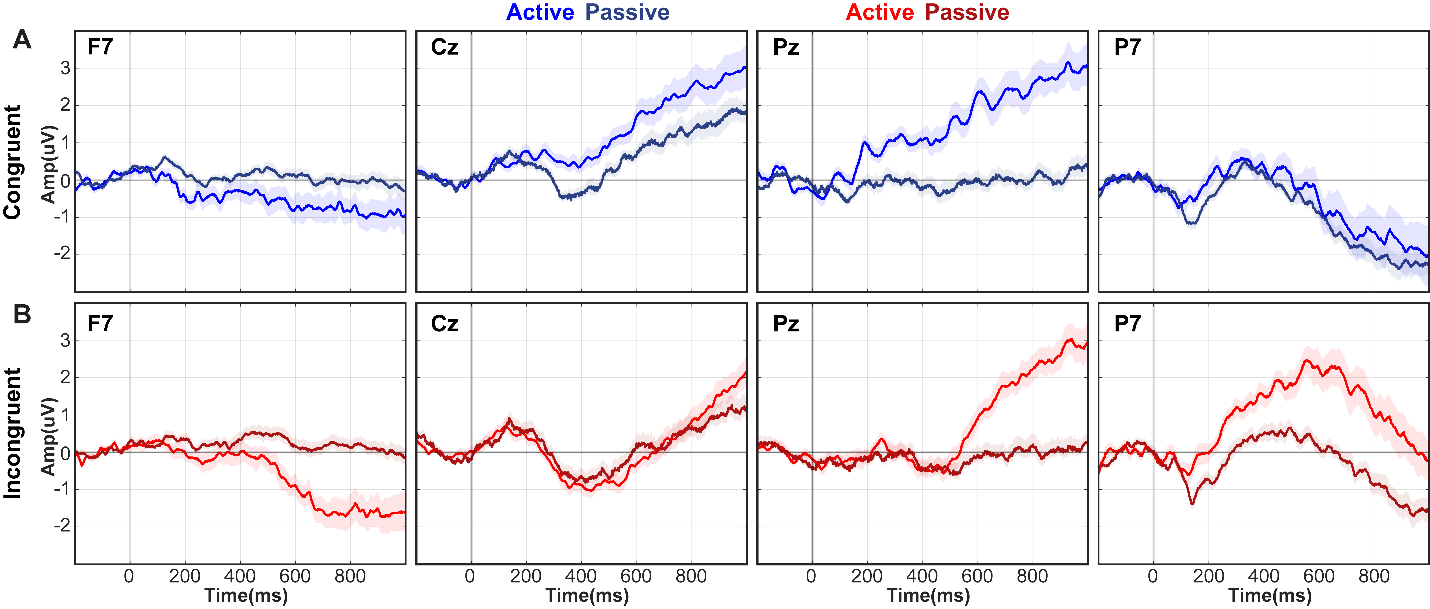

**S1. ERP Plots by Condition.** ERP plots for midline electrodes Cz and Pz, and left parietal electrode P7, show amplitude changes over 1000ms for each paradigm (active vs. passive) in response to each condition (congruent vs. incongruent). Displayed are the responses to the **A.** congruent condition in both active (blue) and passive (navy) paradigms **B.** and incongruent condition in both active (red) and passive (maroon) paradigms. Topographical electrode locations are shown on **Figure 3A**.

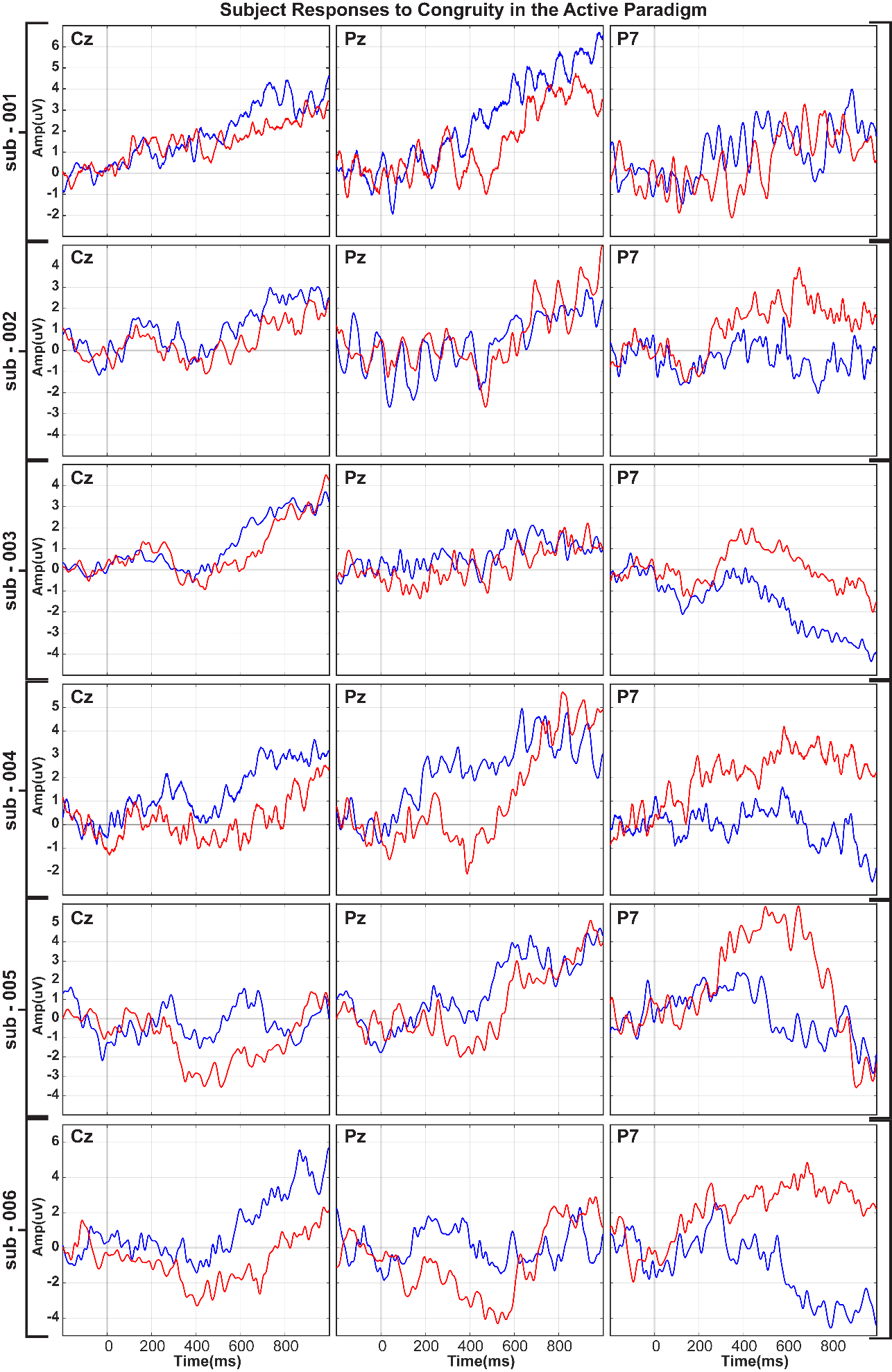

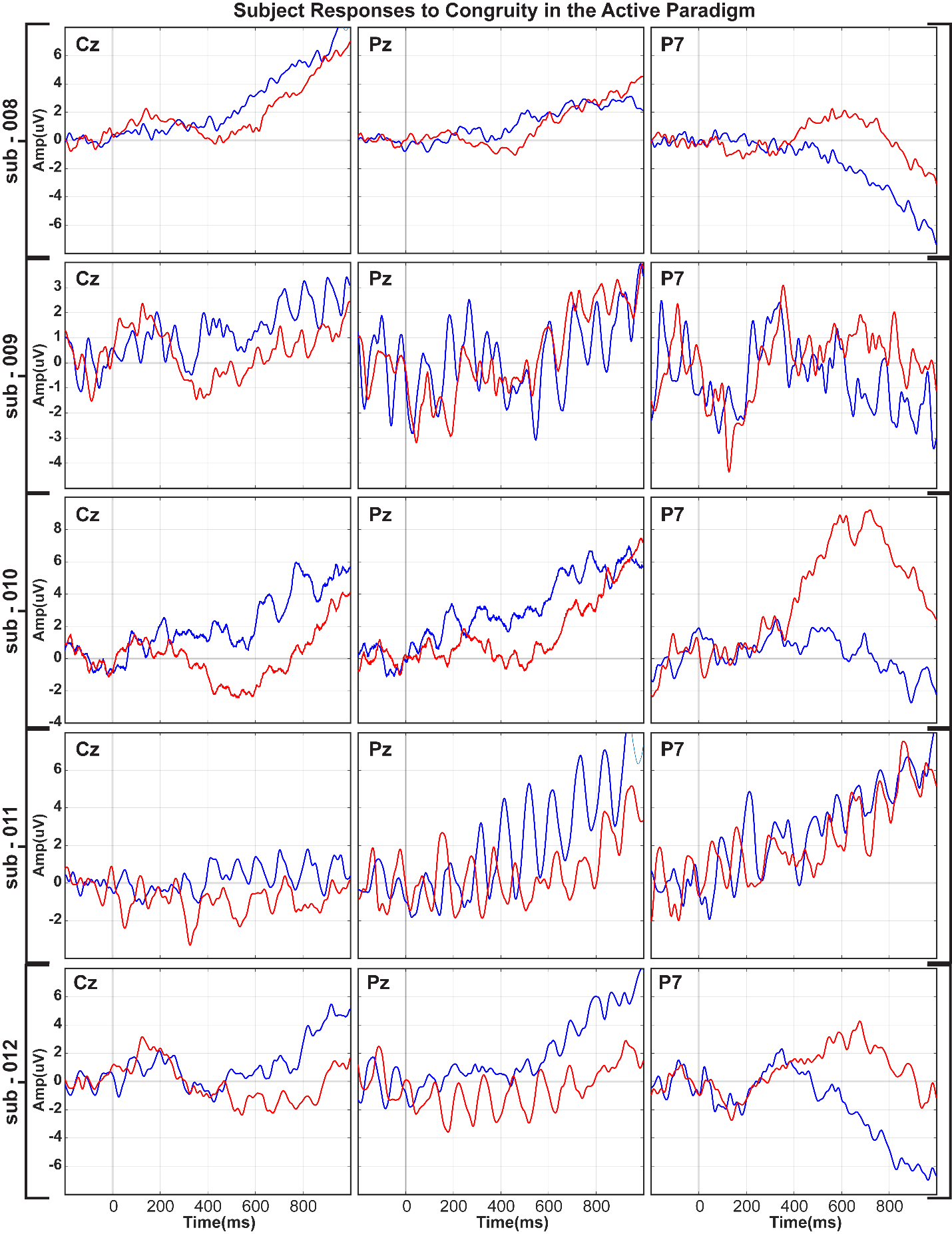

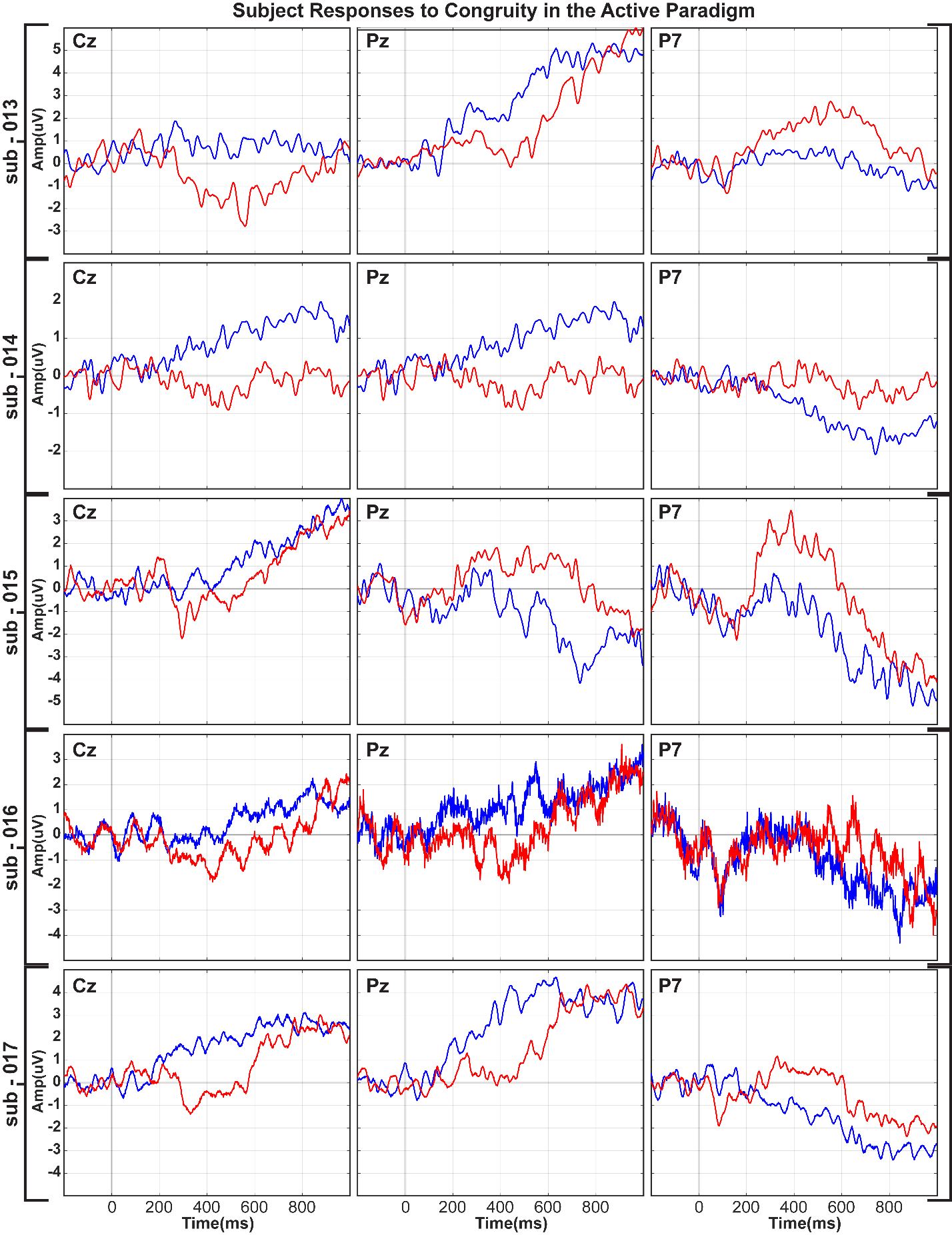

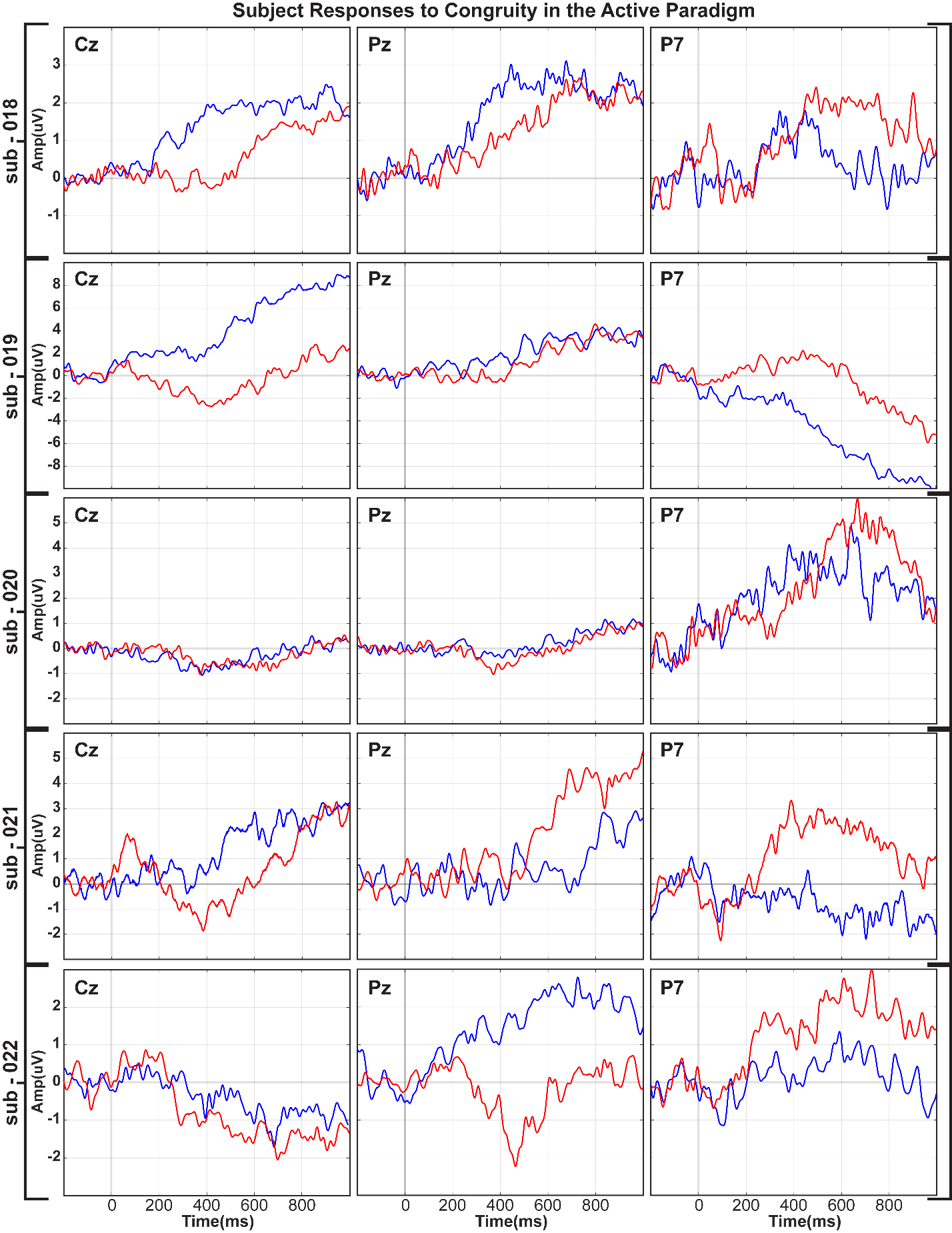

**S2. Individual Subject Responses in the Active Paradigm**

For each individual subject included in the active paradigm, ERP plots for midline electrodes Cz and Pz, and left parietal electrode P7, show amplitude changes over 1000 ms for each condition (Blue, congruent vs. Red, incongruent).

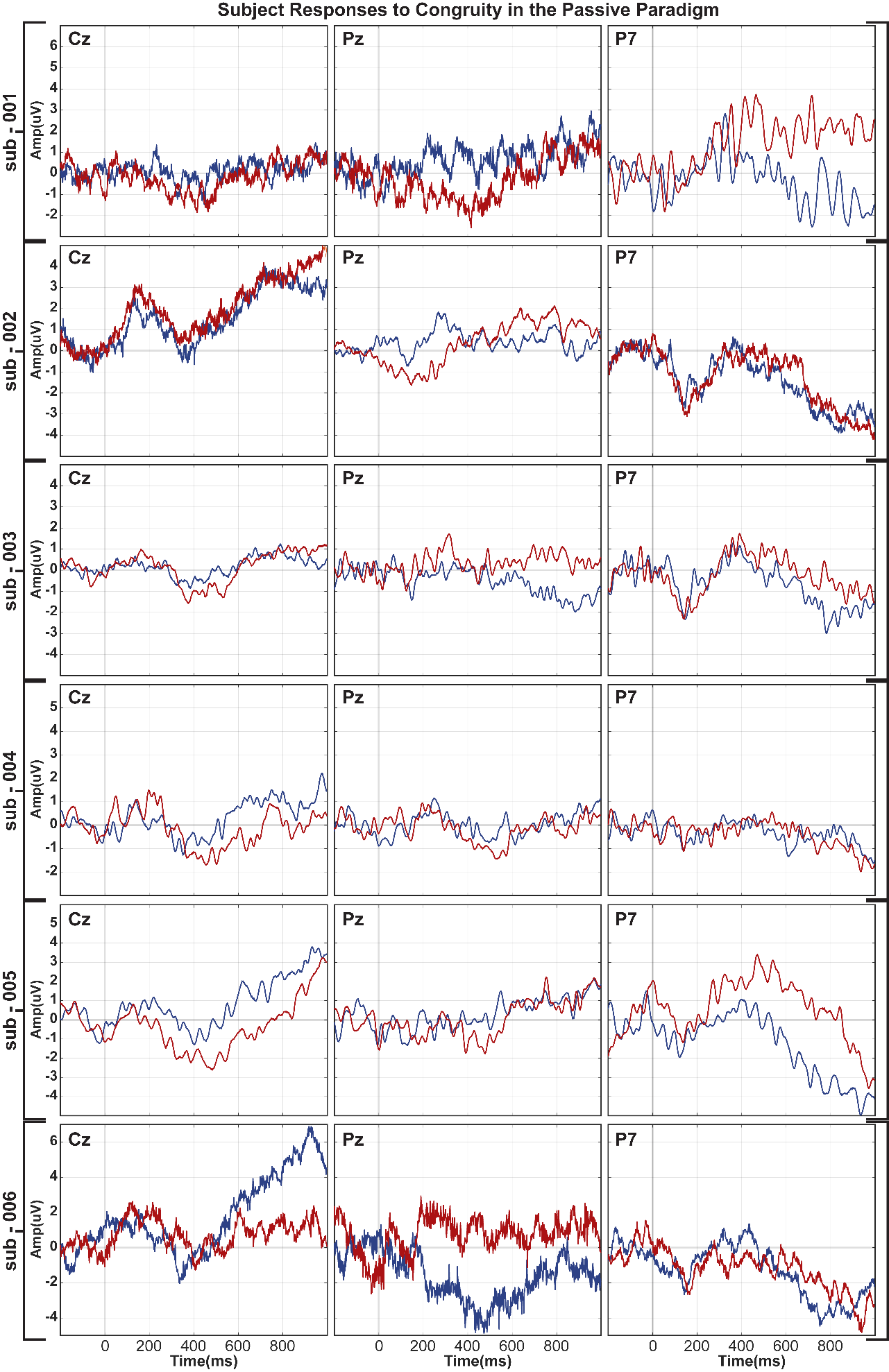

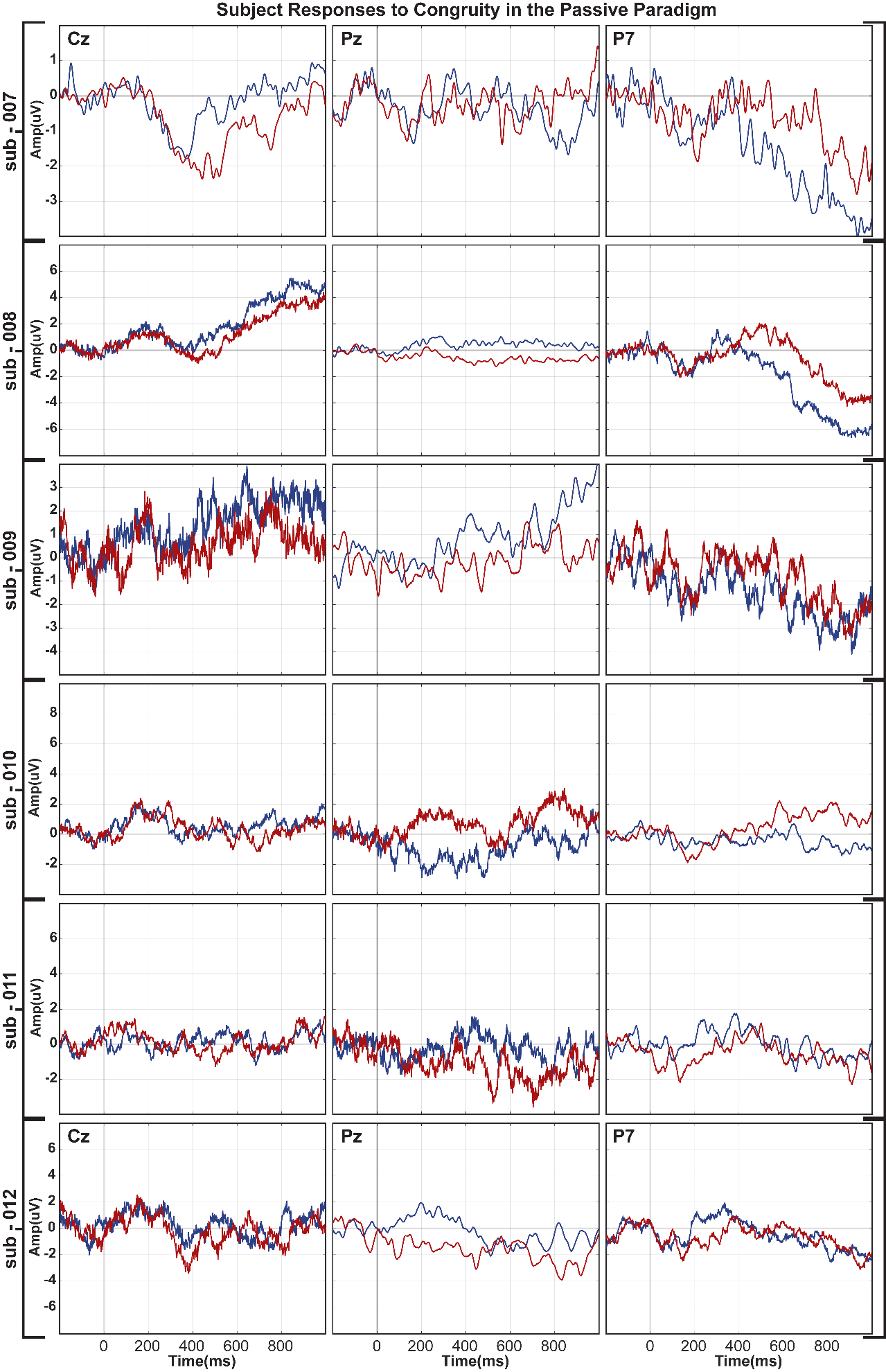

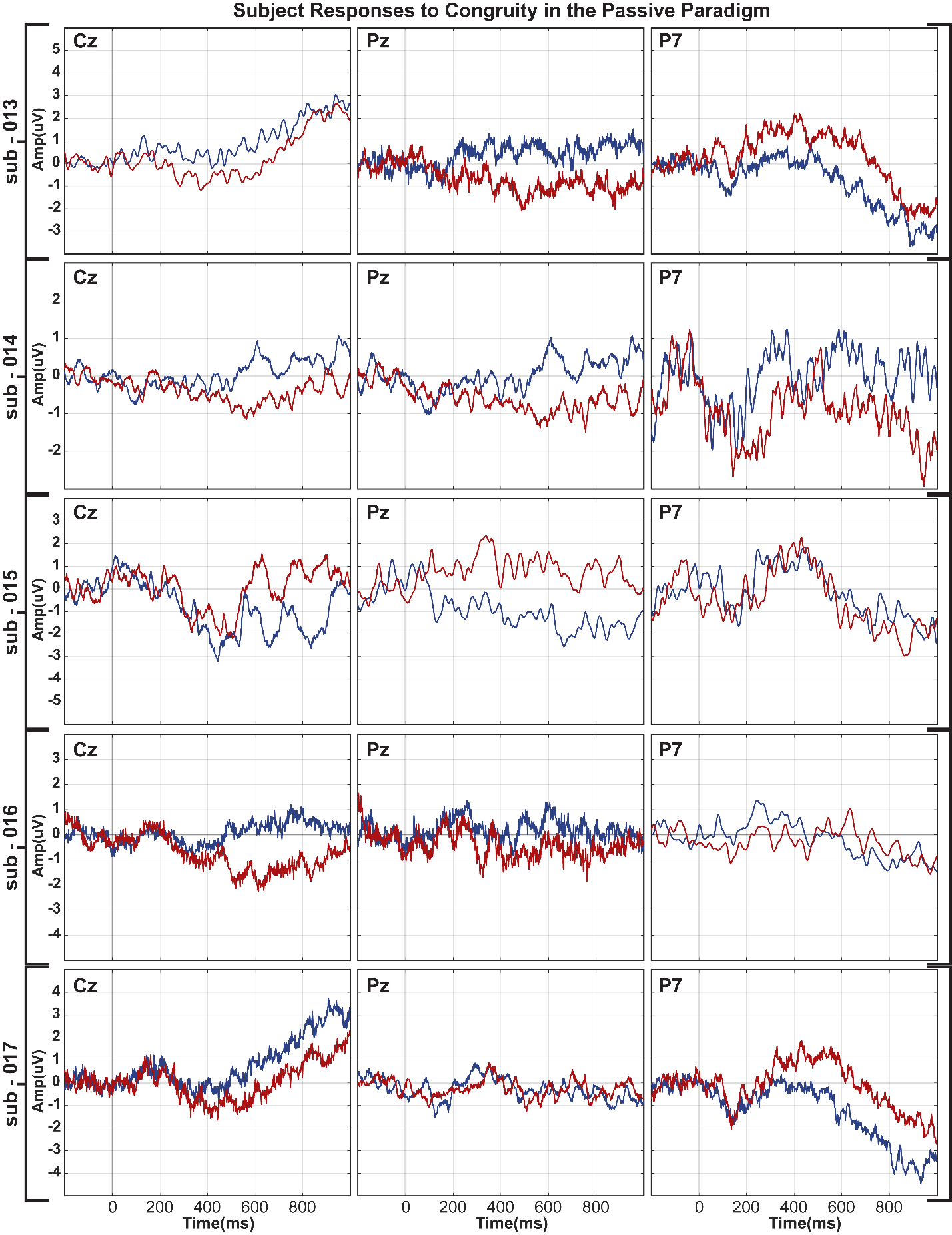

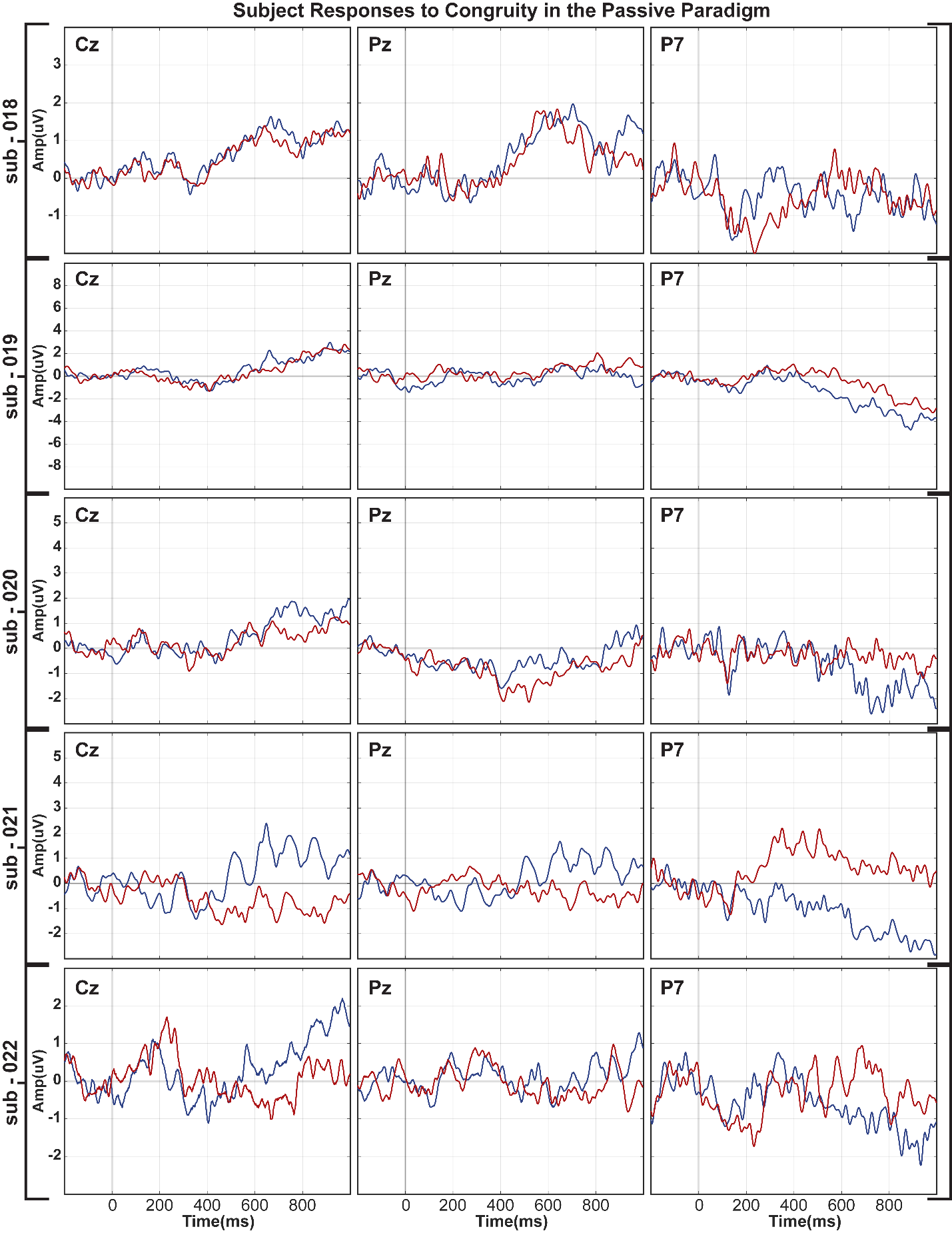

**S3. Individual Subject Responses in the Passive Paradigm**

For each individual subject included in the Passive paradigm, ERP plots for midline electrodes Cz and Pz, and left parietal electrode P7, show amplitude changes over 1000 ms for each condition (Blue, congruent vs. Red, incongruent).
